# Space partitioning by microtubules and molecular motors

**DOI:** 10.64898/2026.11.26.769438

**Authors:** Bhagyanath Suresh, Flora Silberzan, Maxime Fondin, Jérémie Gaillard, Benoit Vianay, Louise Bonnemay, Alexandre Schaeffer, François Nedelec, Alfredo Sciortino, Laurent Blanchoin, Manuel Théry

**Author notes:** these authors contributed equally to the work (Bhagyanath Suresh, Flora Silberzan).

## Abstract

Cells contain radial arrays or polar bundles of microtubules that support their main functions. Active matter made of microtubules and molecular motors can transiently assemble radial asters or form nematic structures that continuously extend and fold. Whether such networks can reach dynamic steady states, as seen in cells, is not known. Here we asked whether confining two oppositely directed motors could generate dynamic, stable and regular organizations. Microtubules alignment and polarity sorting formed linear stacks that partitioned the available space into domains enriched in one motor or the other. Motor ratio, microtubule number and length, and the size and shape of the confinement determined whether the resulting domain organizations were radial, polar, bipolar or multipolar. Similar patterns formed in living cells upon microtubule stabilization. These findings reveal regular, dynamic steady-states in confined active matter and open new ways to understand the diversity of microtubule network architectures in cells.

## Introduction

In cells, microtubules (MTs) serve as tracks for intracellular transport. They are intrinsically oriented by the ordered assembly of α/β-tubulin dimers (*1*, *2*). One end, the so-called “minus-end” terminates with an α-tubulin, the other, the “plus-end” with a β-tubulin. Along the shaft, molecular motors are sensitive to the orientation of these subunits and move specifically toward either the plus or the minus end (*3*, *4*). MTs thereby, guide intracellular transport and direct the spatial compartmentalization of organelles (*5–9*).

The architecture of the network, that is, the position and orientation of MTs, directs the opposite fluxes of motors and thereby determines the spatial organization and polarization of the main compartments, which, depending on cell type, are described as apical and basal poles, front and back, dorsal and ventral sides (*10*, *11*). Distinct cell types come with diverse architectures: radial arrays are common in proliferating cells, whereas differentiated cells rely on various types of polarized arrays made of aligned MTs oriented in the same direction (*12–14*). As cells respond to chemical and physical cues, they can transdifferentiate and switch from radial to polar arrays, or reverse the orientation of their polar arrays (*15–18*). Despite their fundamental role in the regulation of key cellular functions, the mechanisms directing the architecture of MT networks are not yet fully understood.

Because MTs are unstable polymers that do not form spontaneously in the cytoplasm, the network is generally considered to be organized by the regulation of MT dynamics. Nucleating factors are present for example in the centrosome or Golgi apparatus, which can thus initiate MT growth and anchor their minus-ends (*19*, *20*). Molecular complexes, diffusing in the cytoplasm or attached to the cell cortex, regulate the growth and the capture of MT plus-ends (*21*). This raises the question of how these organelles and molecular complexes are positioned. However, in parallel to the regulation of MT dynamics, the network is also organized mechanical forces. Indeed, molecular motors not only travel along MTs but also displace and reorient them, thereby dramatically affecting network architecture (*14*, *22*). The contribution of motors to the spatial organization of MTs is manifest during cell division (*23*). The position and transport of MTs in mitotic and meiotic spindles rely on the opposite forces applied by minus and plus-end oriented motors (*24*). But motors also regulate the position and orientation of MTs in interphase and differentiated cells (*25–30*). However, the physical principles of subcellular self-organization by mechanical forces are still unresolved (*31*), and the actual role of motors in the organization of the broad diversity of arrays formed by MTs in interphase remains to be elucidated.

Indeed, the role of molecular motors in spindle organization was mainly revealed by in vitro reconstitution experiments whereby purified motors were monitored on stabilized MTs to measure their movements and biophysical properties (*32*). Mixing purified motors with stable MTs, a system later termed active matter, led to a variety of architectures offering numerous insights into network self-organization properties (*33*). Complexes made of several motors attached to each other can bridge several MTs simultaneously, and, by moving toward a specific end can cluster them together and form vortices or asters (*34–36*). However, this effect is more effective if the motors are processive, because if a motor detaches before reaching the MT end, it might reattach to another MT of different polarity. In a bundle, MTs of opposite polarities then slide along each other and form extensile bundles (*37–39*). As these bundles extend, MTs become polarity sorted, and if the overlap vanishes, the bundle will cease to extend. However, network density forces these sorted MTs to bend and encounter MTs of opposite polarity, with which they can form another extensile bundle. The network is thus maintained in a continuously extending and folding state, in which a global order is maintained but permanently disrupted locally (*40*, *41*).

Instead of MT-crosslinking motors complexes, immobilized or membrane-bound motors can also move MTs (*42*, *43*) and drive their nematic ordering when present at high density (*44*, *45*). In addition, in response to two types of fixed motors walking in opposite directions, moving MT can be held in place in a compressive tug of war (*46–50*). Recently, we combined these two features and found that if the two types of motors are not fixed but bound to lipid membrane, they can generate concentration gradients at each end of the immobilized MTs as they walk and accumulate at both ends (*51*). These concentration gradients act on adjacent MTs: they attract MTs of same orientation and repel MTs of opposite orientation. This process leads to the assembly of bands of polarity-parallel MTs that act as active barrier between domains containing the plus- and minus-end directed motors. We call “plus-domains” the region enriched in “plus-end motors”, and likewise denote “minus-domains” (*51*) (Fig 1A, Movie S1). The ratio of oppositely directed motors defines the relative areas of the two domains, and their overall appearance can explained by theory of active phase separation and recapitulated in computer simulations (*52*).

**Figure 1:**
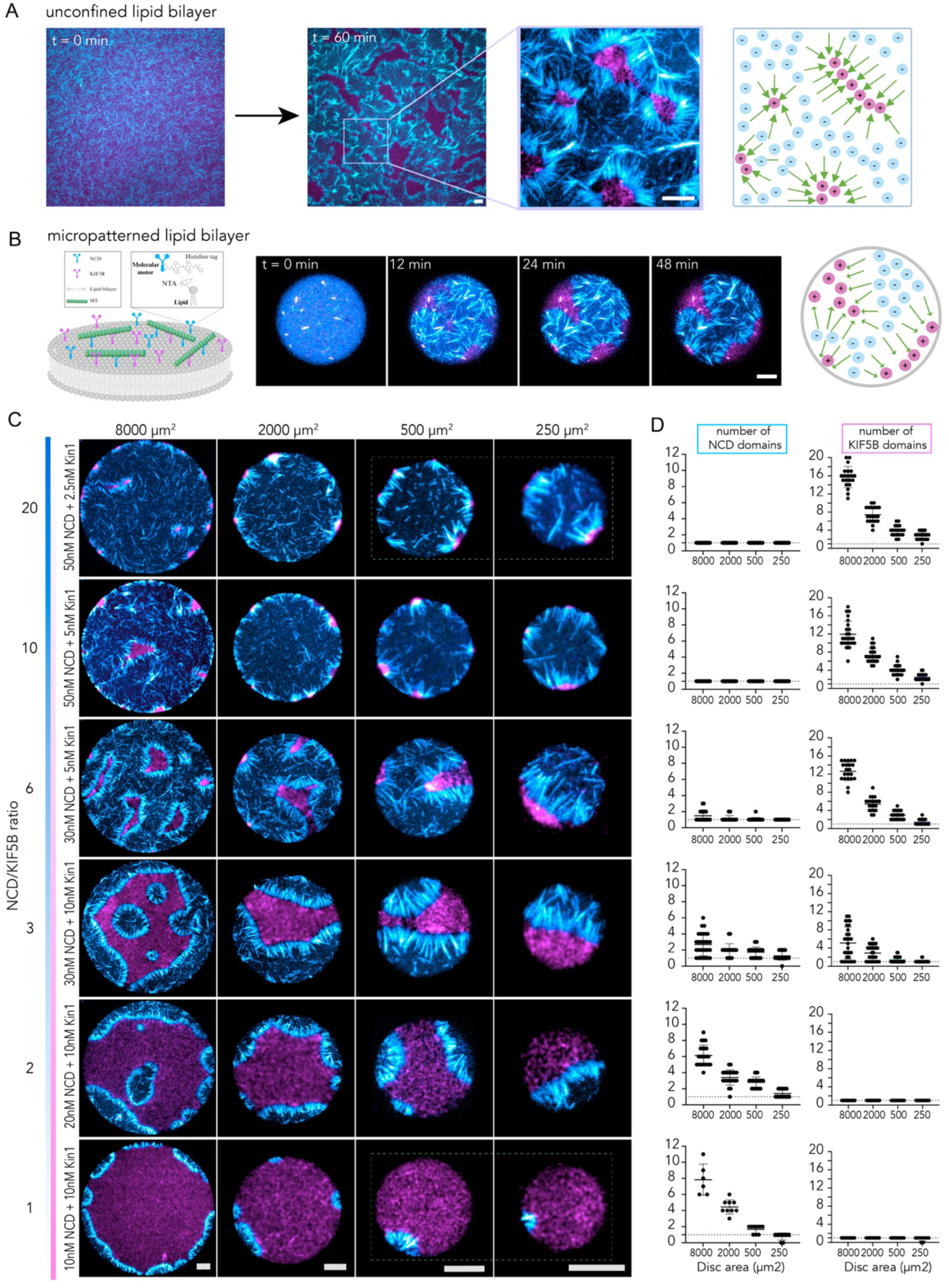
Confining area and motors ratio define the number of partitions. A-Left, fluorescence images showing the self-organization of KIF5B (magenta) and NCD (cyan) on an unconfined supported lipid bilayer (SLB) at 0 and 60 min, with a magnified view of the resulting motor domains. Right, schematic representation of the self-organized motor domains. B-Left, time-lapse images of motor domain formation within a circular micropatterned SLB, acquired at 12-min intervals. Right, schematic representation of the resulting space partitioning. C-Fluorescence images showing motor domain organization within circular confinements of decreasing area (8000, 2000, 500, and 250 µm², left to right) and varying NCD/KIF5B concentration ratios (10/10, 20/10, 30/10, 30/5, 50/5, and 50/2.5 nM, bottom to top). Decreasing confinement area reduces the number of motor domains. At low relative NCD concentrations, a single peripheral NCD domain forms, whereas at low relative KIF5B concentrations, multiple smaller peripheral KIF5B domains persist. D-Quantification of the number of NCD (cyan, left) and KIF5B (magenta, right) domains as a function of confinement area for the motor concentration combinations shown in (C). Data are presented as mean ± SD. Sample sizes for NCD (n = 6, 23, 29, 21, 25, and 21 at 8000 µm²; n = 9, 23, 29, 21, 25, and 21 at 2000, 500, and 250 µm²) and KIF5B (n = 9, 23, 29, 21, 24, and 20 at each confinement area) correspond to NCD/KIF5B concentrations of 10/10, 20/10, 30/10, 30/5, 50/5, and 50/2.5 nM, respectively, where n represents the number of individual circular confinements analyzed. In all panels, scale bars represent 10 µm.

Importantly, physical and geometrical boundaries are known to influence cytoskeleton network self-organization in vitro and in living cells (*53*). Indeed, the confinement of contractile networks of microtubules could lead to their coalescence into a single, large, centered aster or peripheral bundles depending on the confining volume (*54–57*). The confinement of extensile nematic led to the selection of specific wavelength according to channel dimensions and the transition from turbulent into coherent flows (*58–61*). Here we asked whether and how spatial boundary conditions guide the segregation of oppositely directed motors into distinct domains, and whether this process could contribute to the organization of MT networks in cells.

## Results

### Partitioning of confined space

We used maskless photolithography to restrict lipid attachment and diffusion to regions of defined size and shapes (*62*). This forced the motors bound to the diffusing lipids to form their respective domains within these micropatterned regions (Fig. 1B, movie S2). Not surprisingly, the number of plus and minus-domains diminished with the size of the confining area (columns in Fig. 1C, D, movie S3). Because the domains also depend on motor ratio (*51*), we tested several motor ratios for various confining size (rows in Fig 1C,D). In large areas, increasing the relative proportion of minus-end directed motors changed organization from a large central plus-domain with multiple small peripheral minus-domains into an inverted but equivalent configuration, where a minus-domain is surrounded by plus-domains (Fig 1C, left column, movie S4). Interestingly, in small area, close to characteristic length scale separating unconfined domains, the two motors did not produce equivalent patterns (movie S5). At intermediate motor ratios, a single plus-domain and a single minus-domain of equal size formed, leading to a balanced and highly polarized partition of motors (Fig 1C, right column, middle rows). At low amounts of minus-end motors, the size of the minus-domain was reduced to a single small peripheral spot (Fig 1C right column, bottom, movie S6). In contrast, at low amounts of plus-end motors, multiple small plus-domains formed on the outline (Fig 1C right column, top, movie S7). Thus, one state broke symmetry and appeared polarized as a single peripheral minus-domain formed, whereas the other kept the circular symmetry of the pattern with multiple peripheral plus-domains, as shown by the measurement of the average position of all domains (Figure 2A, B). This difference is likely due to the unequal biophysical properties of the two motors. We reasoned that a weaker motor (NCD) would have to form larger domains because more individual motors would be needed to produce a force opposing the stronger motor (KIF5B), as the forces on MTs are necessarily balanced at the interface between two domains in a steady state. This hypothesis was supported by numerical simulations showing that motors of different strengths were sufficient to recapitulate all these observations (Fig. S1, movie S8).

**Figure 2.**
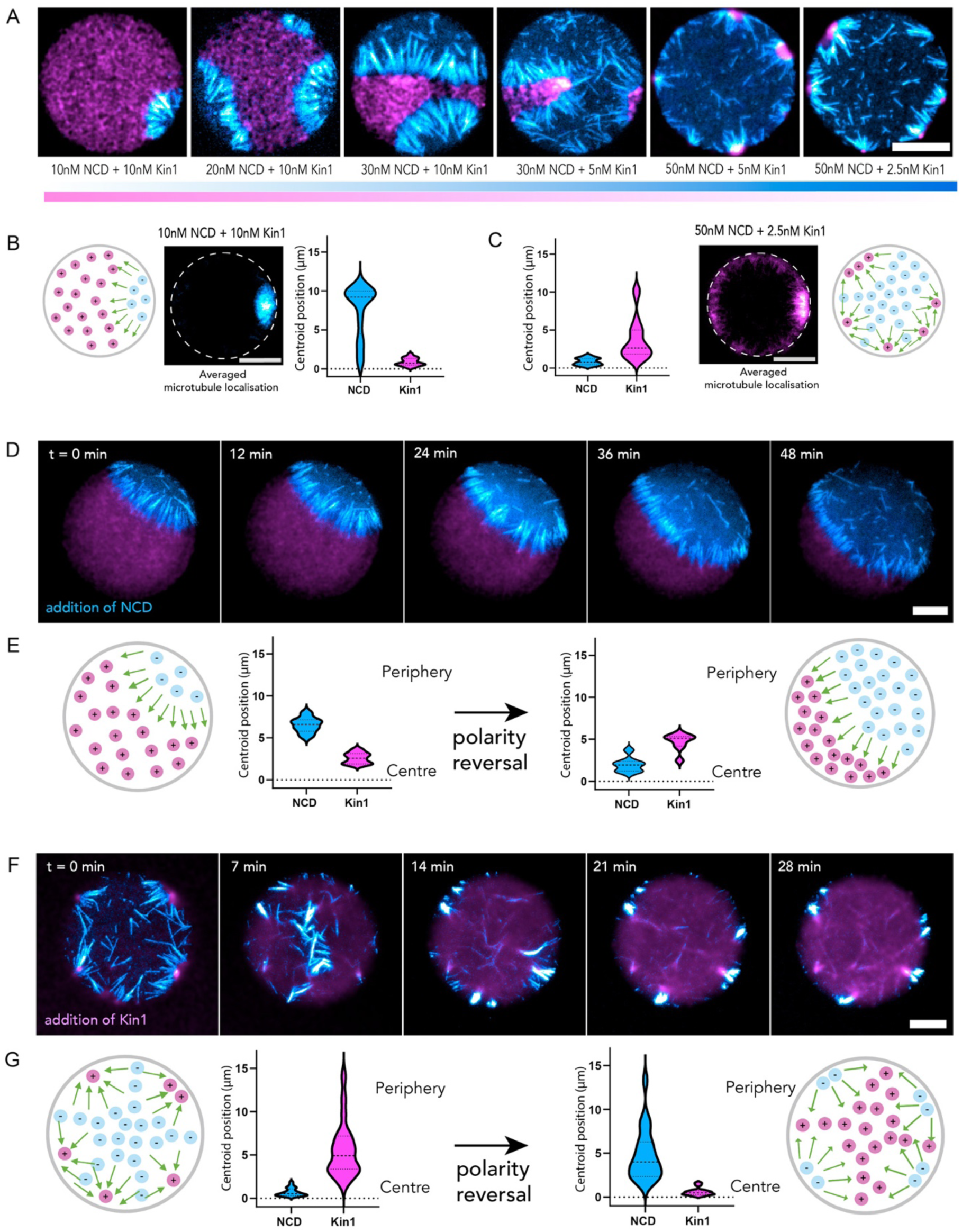
Partitions are responsive to changes in motors concentration. A-Fluorescence images of KIF5B (magenta) and NCD (cyan) self-organizing within circular confinements of 500 µm² at varying NCD/KIF5B concentrations (10/10, 20/10, 30/10, 30/5, 50/5, and 50/2.5 nM, left to right). Increasing the relative concentration of NCD changes the organization from a single peripheral NCD domain to multiple peripheral KIF5B domains. B,C-Schematic representations of the polar (B) and radially symmetric (C) organizations formed at low relative concentrations of NCD and KIF5B, respectively. Averaged localization maps were obtained by thresholding and aligning microtubule images from individual circular confinements. Violin plots show the distribution of the distance of NCD and KIF5B domain centroids from the confinement center (n = 43 for both motors in B; n = 40 for NCD and n = 27 for KIF5B in C). D-Time-lapse images showing the reversal of a pre-existing polar organization following the addition of NCD, acquired at 12-min intervals. E-Schematic representations and quantification of NCD and KIF5B domain centroid positions before and after NCD addition, showing a reversal of their peripheral and central localization. F-Time-lapse images showing the reversal of a pre-existing radial organization following the addition of KIF5B, acquired at 7-min intervals. G-Schematic representations and quantification of NCD and KIF5B domain centroid positions before and after KIF5B addition (before: n = 22 for NCD and n = 21 for KIF5B; after: n = 21 for both motors), showing an inversion of the radial organization. Each violin plot represents the distribution of individual motor domain centroid positions relative to the confinement center. Solid lines indicate the median, and dotted lines indicate the 25th and 75th percentiles. In all panels, scale bars represent 10 µm.

### Polarity reversal

As cells polarize or depolarize - for example, during transitions between epithelial and mesenchymal states (*15*, *16*) - they switch between radially symmetric and polarized MT arrays. Because opposite motor ratios led to radial or polarized arrays in our minimal system, we asked whether we could induce such a transition by changing the motor ratio during the experiment. However, adding motors of one type to a state established by the other motor did not produced the state corresponding to the added motor type, but changed it to another, new, state. Adding minus-end motors to a polarized state formed with an excess of plus-end motors reversed the orientation of the polarized state (Fig. 2D, E, movie S9). Adding plus-end motors to a radial state formed with an excess of minus-end motors reversed the radial state from minus-end in to minus-end out (Fig. 2F, G, movie S10). Thus, instead of switching from a radial to a polarized state, the delayed modification of the motor ratio reversed the orientation of the initial state. By adding the motors sequentially, we obtained configurations not observed when they were mixed from the outset at any ratio, including a single polar plus-domain, or multiple peripheral minus domains. This showed that continuous motor transport along MTs allows the partition to reorganize if motor concentrations are changed over time, and that the steady state depends not only on the motor ratio but also on the system’s history.

### Partition depends on MT number and length

Segregated domains are maintained by the active transport of motors, and the bands of polarity-parallel MTs act as active barrier by directing each motor into its respective domain. Intuitively, to be effective this barrier requires a minimal density of MTs, otherwise, motors would diffuse between adjacent MTs, creating a leak that would destabilize the segregated domains. This critical density of MTs likely sets the length of the interface that can be assembled with a given number of MTs. To test this idea, we varied the number of MTs on a given area loaded with a given ratio of motors (Fig. 3A). As expected, more MTs formed longer interfaces and thereby partitioned the space into more domains (Fig. 3B, movie S11). However, on small area (500 µm^2^), the number of domains increased less rapidly with the number of MTs, because their formation appeared to be limited by the length of the MTs separating the domains (Fig. 3C and D, movie S12). Indeed, one can expect that the distance between domain centers cannot be smaller than the MT length. This further suggested that MT length could also be a key regulator of domain organization. Preliminary tests on non-confined systems revealed that domains disassemble when elongating MTs contacted the opposite interface (Fig. S2A, movie S13). This can be seen as the outcome of force imbalance. Indeed, MTs remain steady in place only as long as the forces generated by the opposite motors balance (*51*). If MTs elongate enough to overlap with a third domain, one motor wins, and the MT is pushed away from the interface (Fig. S2B). We tested the role of MT length under confinement by preassembling MTs and motors and subsequently adding free tubulin. Without tubulin addition, pre-existing domains remained steady (Fig. S2C, movie S14). The MT elongation resulting from addition of free tubulin led to the disassembly of the smallest domains. On large patches (2000 µm^2^) MT lengthening reorganized multipolar architectures into bipolar networks, made of a single central plus-domain and two peripheral minus-domains (Fig. 3 D, E, movie S15). On smaller patches (500 µm^2^), MT elongation from the same initial biochemical conditions led to polar networks, made of a single plus-domain and a single minus-domain separated by a band of parallel MTs (Fig. 3 F, G and H movie S16). Hence, the number of motor domains increased with the number of MTs but decreased with their length. The balance of these opposite contributions led to the robust formation of polar or bipolar structures depending on the confinement area and the system’s history.

**Figure 3.**
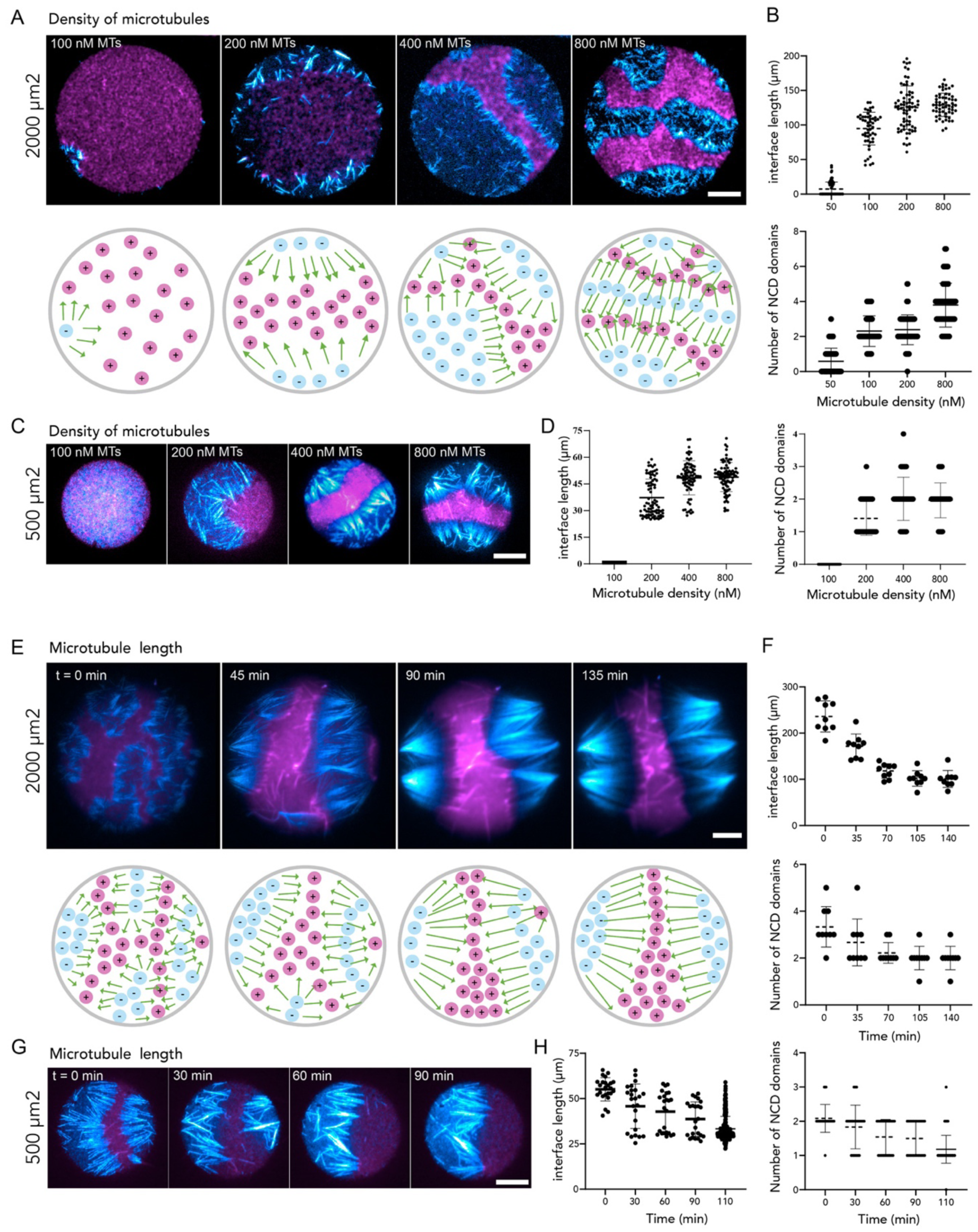
Microtubule density and length regulate the number of partitions. A-Top, fluorescence images showing the self-organization of KIF5B (magenta) and NCD (cyan) within circular confinements of 2000 µm² at increasing microtubule concentrations (50, 200, 400, and 800 nM). Bottom, schematic representations of the resulting motor domain organization. B-Quantification of the total interface length between motor domains (top) and the number of NCD domains (bottom) as a function of microtubule concentration for the conditions shown in (A). Sample sizes for interface length (n = 84, 52, 70, and 58) and NCD domain number (n = 84, 52, 80, and 59) correspond to microtubule concentrations of 50, 200, 400, and 800 nM, respectively. C-Fluorescence images showing the organization of KIF5B and NCD domains within circular confinements of 500 µm² at increasing microtubule concentrations (100, 200, 400, and 800 nM). D-Quantification of the total interface length between motor domains (left) and the number of NCD domains (right) as a function of microtubule concentration for the conditions shown in (C). Sample sizes for interface length and NCD domain number (n = 31, 81, 76, and 81) correspond to microtubule concentrations of 100, 200, 400, and 800 nM, respectively. E-Top, time-lapse images showing the reorganization of motor domains following the addition of free tubulin to promote microtubule elongation within circular confinements of 2000 µm². Bottom, schematic representations of the resulting transition from a multipolar to a bipolar organization. F-Quantification of the total interface length between motor domains (top) and the number of NCD domains (bottom) during microtubule elongation (n = 9 confinements at each time point). G-Time-lapse images showing the reorganization of motor domains following microtubule elongation within circular confinements of 500 µm², resulting in a polar organization. H-Quantification of the total interface length between motor domains (top) and the number of NCD domains (bottom) over time for the conditions shown in (G). Increasing microtubule density promotes the formation of longer interfaces and additional motor domains, whereas microtubule elongation reduces interface length and domain number. Each point represents an individual circular confinement; horizontal lines and error bars represent the mean ± SD. In all panels, scale bars represent 10 µm.

### Partition depends on confinement shape

We then tested whether the shape of the confinement could affect the formation of motor domains. A mixture of 400 nM MTs, 10 nM KIF5B and 10 nM NCD was introduced onto micropatterns with circular and elliptical shapes, varying aspect ratios at equal areas. In these biochemical conditions, on small areas (500 µm^2^), minus motors (NCD) tended to form preferentially a single domain on circular shapes, whereas elongated shapes increased this number to two minus-domains, at the tips of the ellipse (Fig. 4A, B, movie S17). On larger areas (2000 µm^2^), more minus-domains formed on circular shapes, whereas elliptical shapes reduced this number to two (Fig. 4A, B, movie S18). Thus, the formation of bipolar organization was strongly enhanced on elliptical shapes. We again interpreted this to be a consequence of difference in motor’s properties: high local curvature allows a larger number of weaker NCD motors to be enclosed behind a shorter interface to oppose the stronger KIF5B motors.

**Figure 4.**
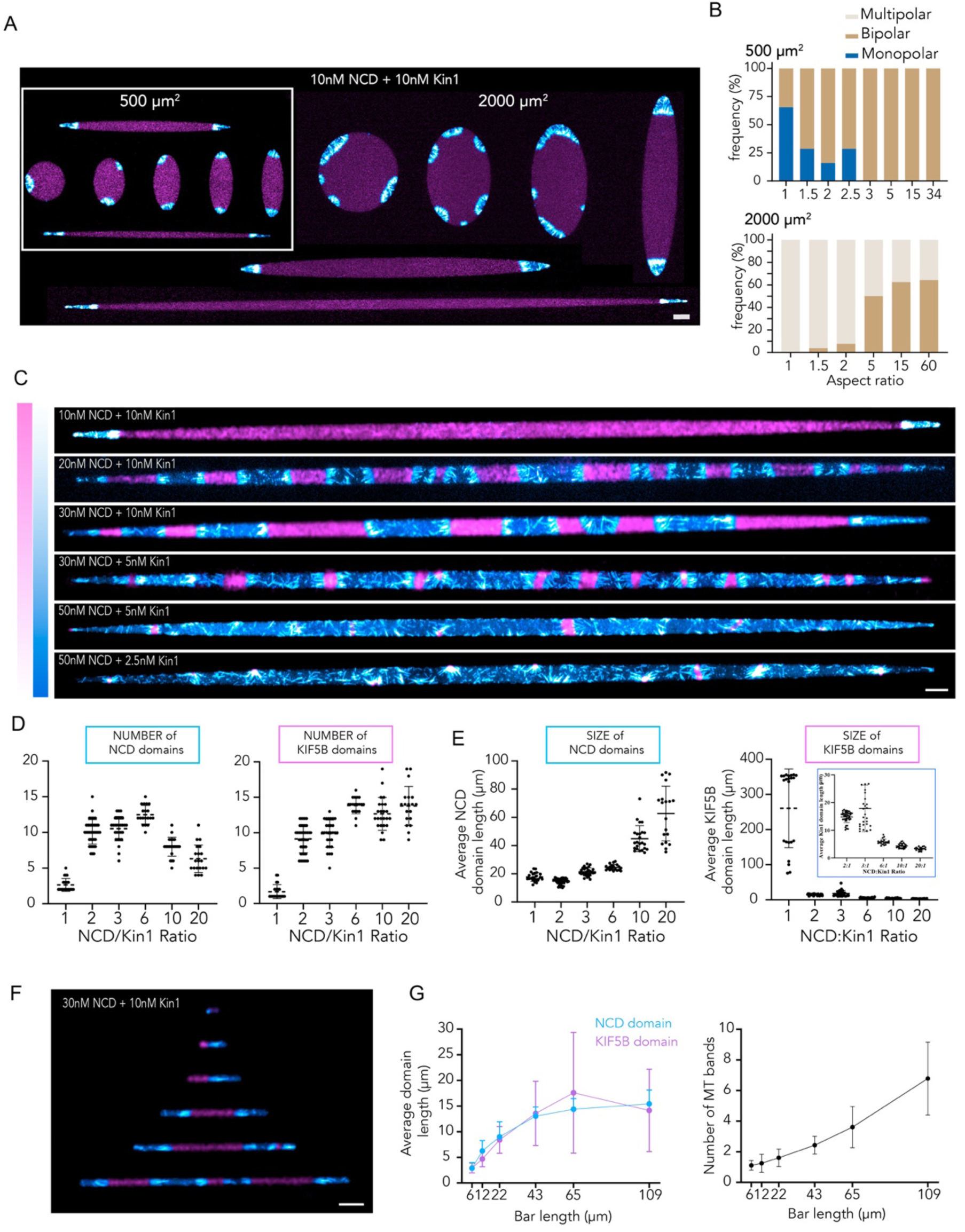
Confinement geometry and motors ratio direct the position and number of partitions. A-Fluorescence images showing the self-organization of KIF5B (magenta) and NCD (cyan) within elliptical confinements of increasing aspect ratio (starting with circles) at fixed areas of 500 and 2000 µm², with 10 nM NCD and 10 nM KIF5B. Increasing confinement elongation promotes the formation of bipolar motor-domain organization. B-Quantification of the relative frequency of monopolar (blue), bipolar (brown), and multipolar (gray) organizations as a function of confinement ellipticity for areas of 500 µm² (top) and 2000 µm² (bottom). Confinements were classified according to the number of NCD domains formed (one, two, or more than two domains, respectively). Sample sizes for 500 µm² confinements (n = 26, 7, 25, 7, 7, 19, 26, and 7) correspond to ellipticities of 1, 1.5, 2, 2.5, 3, 5, 15, and 34, respectively. Sample sizes for 2000 µm² confinements (n = 26, 26, 26, 24, 24, and 28) correspond to ellipticities of 1, 1.5, 2, 5, 15, and 60, respectively. C-Fluorescence images showing motor domain organization within highly elongated confinements of 2000 µm² at increasing NCD/KIF5B concentrations (10/10, 20/10, 30/10, 30/5, 50/5, and 50/2.5 nM, top to bottom). Increasing the relative concentration of NCD leads to the formation of alternating motor domains, followed by an increase in the size of NCD domains at higher ratios. D-Quantification of the number of NCD (cyan, left) and KIF5B (magenta, right) domains as a function of motor concentration ratio for the conditions shown in (C). Sample sizes for NCD (n = 22, 40, 29, 19, 24, and 20) and KIF5B (n = 22, 40, 29, 19, 24, and 19) correspond to NCD/KIF5B ratios of 1:1, 2:1, 3:1, 6:1, 10:1, and 20:1, respectively. E-Quantification of the average length of NCD (cyan, left) and KIF5B (magenta, right) domains for the conditions shown in (C). Sample sizes for NCD (n = 22, 40, 29, 19, 24, and 19) and KIF5B (n = 22, 40, 28, 19, 24, and 19) correspond to the same motor concentration ratios. F-Fluorescence images showing the organization of KIF5B (magenta) and NCD (cyan) within elongated confinements of increasing length (6, 12, 22, 43, 65, and 109 µm) at fixed motor concentrations of 30 nM NCD and 10 nM KIF5B. G-Quantification of the average NCD (cyan) and KIF5B (magenta) domain lengths (top) and the number of microtubule bands (bottom) as a function of confinement length (n = 28 confinements for each length). Domain lengths approach a characteristic size, whereas the number of bands increases with confinement length. For the scatter plots in (D), (E), and (G), each point represents an individual confinement; horizontal lines and error bars represent the mean ± SD. In all the panels, scale bars represent 10 µm.

We noticed that the variability of domains formation was lower on very elongated ellipses. We therefore used these nearly linear shapes to revisit the contribution of the motor ratio under these more reliable conditions. As we increased the concentration of minus-end motors from 10 to 20 nM, the single central plus-domain fragmented into alternating minus and plus-domains (Fig. 4C top rows, Fig. 4D,E, movie S19). MTs formed regularly spaced bands perpendicular to the long axis of the ellipse, at the frontier between domains. As minus-end motors were added further, the size of the minus-domains became larger and fewer, eventually down to a single central minus-domain (Fig. 4C, bottom row). The number and size of plus-domains followed the opposite trend (Fig. 4, D and E) because the total confining area was constant. Thus, the respective sizes of plus and minus-domains accurately reflected the ratio of the corresponding motors.

Interestingly, we found that above a critical pattern length, corresponding to the domain size set by the biochemistry (15 µm for a NCD:KIF5B ratio of 3:1), the size of domains no longer grew (Fig. 4, F and G). When the patterns were made longer, the number of domains increased proportionally (Fig. 4, F and G, movie S20). This linear increase of domain number with pattern length was robust to variations in the motor ratios (Fig. S3), confirming that the biochemistry sets the size of the domains as expected from theory (*43*, *44*).

### Partitioning of intra-cellular space

Do these partitioning rules apply to MT organization in cells? In most cells, MTs are constantly growing and shrinking with an average lifetime of few minutes. Under such conditions, their position is thus foremost determined by their nucleation site. If motors reorganize MTs, they have only a few minutes to do so before MTs disassemble. Thus, the longer the MT persist, the greater the contribution of the motors. We therefore decided to stabilize MTs completely, to inhibit MT nucleation and fully reveal the potential organizing capacities of motors (*63*). PtK2 cells stably expressing GFP-tubulin were treated with 10 µM Taxol to protect MTs from disassembly (*64*). As expected from early studies (*65*, *66*), the initial radial organization of MTs was not maintained and, within 5 to 10 hours, MTs aligned into wide bundles (Fig. 5A, movie S21). This showed that an organizing mechanism acts in parallel with MT dynamics and could possibly rely on the sorting of motors of opposite polarity. However, in these conditions, the development of the organizing mechanism was biased by the initial architecture of the network. To better understand the underlying process, we sought to initiate it from a homogeneous state. To that end, MTs were first depolymerized with nocodazole overnight, so that the nucleators could be completely dispersed. Nocodazole was then washed out and replaced by Taxol, so that newly growing MTs would be rapidly stabilized at a short length throughout the cell. In less than 2 hours adjacent MTs aligned and stacked into regularly spaced bands (Fig. 5B, movie S22). The shape, spacing and parallel alignment of the bands were reminiscent of the boundaries we observed between segregated motors in our reconstitution assay.

**Figure 5.**
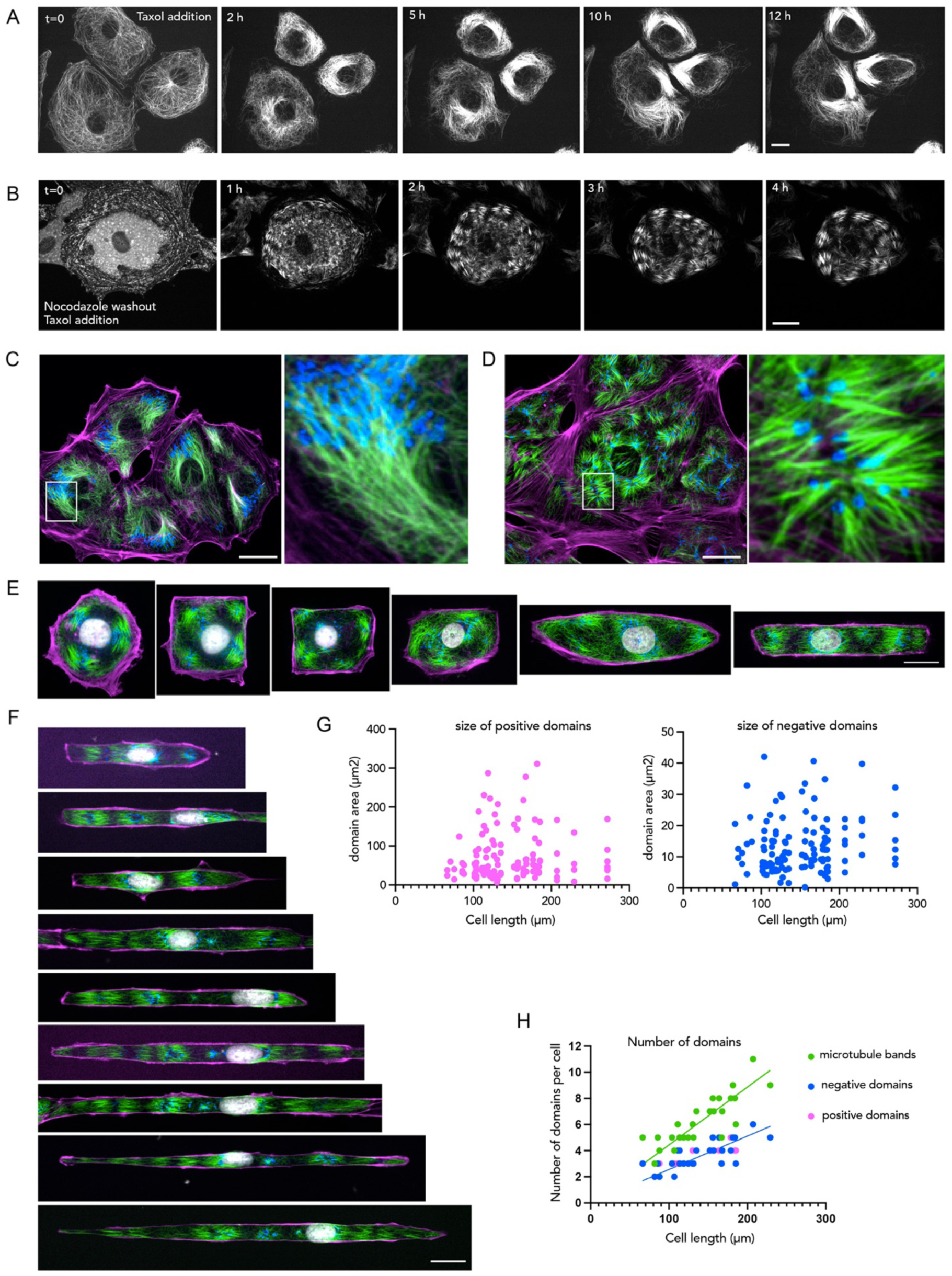
Microtubule stabilization induces cell partitioning in polar domains. A-Time-lapse imaging of a PtK2 cell expressing GFP-tubulin incubated with 10 µM Taxol. Scale bar represents 20 µm. B-Time-lapse imaging of a PtK2 cell expressing GFP-tubulin incubated with 5 µM Taxol after an overnight pre-treatment with 10 µM Nocodazole. Scale bar represents to 20 µm. C, D, E, F-Immunofluorescence images showing actin (magenta), microtubules (green), Golgi apparatus (blue) and nucleus (gray) in fixed PtK2 cells. In (C), cells were treated with 5µM Taxol for 6 hours. In (D), cells were pretreated treated with 10µM Nocodazole overnight and then fixed 1h after Nocodazole washout and addition of 5µM Taxol. In (E and F), cells were pretreated treated with 10µM Nocodazole for 4h and then fixed 3h after Nocodazole washout and addition of 5µM Taxol. In (E and F), cells were plated on 2000 µm² adhesive micropatterns of various aspect ratio and 12 µm-wide lines respectively. Scale bar represents 20 µm. G-Length of microtubule bands as a function of cell length for the experiment shown in panel (F) H-Number of microtubule bands as a function of cell length for the experiment shown in panel (F).

We further assessed the polarity of MTs in these bands by staining for the Golgi apparatus, which is transported to MTs minus-ends by dynein (*67–69*). In cells treated directly with Taxol (Fig 5C) or pretreated with nocodazole (Fig. 5D), Golgi mini-stacks were located on one side of the bands, showing that MTs within a band shared the same orientation. The Golgi stacks were present in one of every two domains. This suggested that the bands separated domains alternately enriched in plus-end- and minus-end-directed motors. However, the bands were not regularly spaced throughout the cytoplasm. To reduce this variability, cells were plated on fibronectin-coated micropatterns to impose a controlled shape. In round cells, the bands and separating domains tended to align with the cell radius (Fig. 5E). However, these conditions were not sufficiently constraining to normalize the shape and spacing of the bands (Fig. S4A). As cell shape were made more elongated, the radial organization progressively switched to a parallel organization, in which bands aligned perpendicular to the long cell axis of the cell. Beyond a width-to-length ratio of 4, the parallel organization was quite reproducible (Sup. Fig. S4B, movie S23). Interestingly, on thin lines of 12 µm, the width of cells was imposed but they could adapt their length depending on their size, which varied up to twofold, likely as a consequence of cell cycle progression (*70*) (Fig. 5F). This allowed us to test the effect of cell length on their intracellular partitioning. We outlined the MT bands and the domain separating them, which were termed positive or negative depending on the presence or absence of Golgi apparatus. The size of the domains, whether positive or negative appeared independent of cell length (Fig. 5G). Negative domains were smaller than positive domains (13+/-5 and 77+/-33µm^2^ respectively). Consistently, the number of domains, and therefore of separating MT bands, increased linearly with cell length (Fig. 5H, Sup. Fig S4C). This showed that not only the shape and organization of MT bands and separating domains were similar to those formed by polarity sorting of MTs and opposite motors in our minimal assay, but also that their variation with respect to geometrical constraints was the similar. This further suggested that stable MTs were actually organized by motor-based polarity sorting.

## Discussion

Altogether our result show that the combined actions of motors of opposite polarity can drive active matter toward a range of organizations, extending the previous description of collective motion, contractile asters and extensile nematics. Their ability to generate spatial patterns by active phase separation leads, under confinement, to a finely tuned and deterministic process of space partitioning. The ratio of plus-end and minus-end directed motors sets the relative sizes of the partitions, and the number and length of MTs set the length of the interface and the number of partitions. The peripheral boundaries, which in this study act as physical barrier for motors but not for MTs, influence the position of the motor partitions and the orientations of the interfaces. Because these interfaces are maintained by the permanent transport of motors in opposite directions along MTs, the initial partition can be reconfigured in real time if motor concentrations or MT length or density are changed. Interestingly, as previously observed in the assembly of MT asters in egg extracts (*71*, *72*), the transformation of initial states in response to biochemical changes may not follow a reversible path and instead display hysteresis. Indeed, certain states—such as one domain of strong motors or many domains enriched in weak motors—could be reached only if the components are added in a specific temporal sequence; they were not observed when all components were mixed at the outset in any ratio. Thus, the final partitioning depends on both the biochemical conditions and the system’s history.

Early observations of contractile asters in mixture of MTs and motors revealed how MT ends can be clustered in the absence of nucleating and anchoring centers, and similar mechanisms are present in cells in interphase (*73*) and at the poles of mitotic spindles (*74*). The potential role of extensile nematic in cells is less clear but the process is reminiscent of MT sliding in growing neuronal protrusions (*14*, *28*, *75*, *76*). These mechanisms are limited by the duration of motor transport along MTs, which is increased by stabilizing the MTs. We found that the stabilization of MTs leads to space partitioning in cells and that the size and position of the partitions follow rules similar to those observed in our in vitro experiments. In both cases, the number of partitions increases linearly with compartment size, suggesting that both rely on the same organizing mechanism. Our in vitro results further show that depending on the ratio of opposite motors and the length and number of MTs, their partitioning can lead to radially symmetric (plus-ends in or out), polar, bipolar or multipolar states. These observations may help us better understand living cells, as many reported *in vivo* systems share similar features. Indeed, mitotic and meiotic spindles can be unstable and generate multipolar arrangement, whose bipolarity can be restored by molecular motors independently of MT nucleating and anchoring centers (*77–79*). Coenocytes and syncytia, which contain several nuclei following multiple rounds of endoreplication or cell fusion, can adopt segmented or unified partitioning. As myoblasts fuse to form multinucleated myotubes they transit from a focused and radial network to linear network organization (*80*, *81*). The polarity sorting of MTs by motors first support cell integration and nuclei clustering (*82*), then cell segmentation and nuclei separation (*83–85*). Similar variations in motor-based partitioning can be observed in other multinucleated cell types such as megakaryocytes (*86*), osteoclasts (*87*), fungi Hyphae (*88*, *89*) and animal early syncytial embryos (*90*, *91*). Conversely, MT stabilization and motor-based transport and polarity-sorting turn the multicellular partitioning of the germline cyst into an integrated and polarized network pointing toward of a single oocyte (*92*). However, the respective roles of MT nucleating complexes and molecular motors overlap and can hardly be disentangled (*73*, *92*, *93*). Local regulations of MT growth and shrinkage are generally better characterized (*12*), but our results show that plus- and minus-end directed motors together are sufficient to generate this diversity of partitioning patterns. Further works should elucidate how the two organizing processes compete or synergize to establish cell symmetry and polarization.

## Acknowledgements

This work was supported by fundings from the Agence Nationale pour la Recherche (grants No AAPG2022-PRC-SHARP to M.T., ANR-24-CE13-7996-MICRON to M.T., Chaire d’excellence en Bio-Santé du programme France 2030 ANR-23-CHBS-0013 to M.T.) and from the European Research Council under the European Union’s Horizon 2020 research and innovation programme (grant No 951430 to F.N.). AS has received funding from the European Union’s Horizon 2020 research and innovation programme under the Marie Sklodowska-Curie grant agreement No101108326.

## Author contributions

Conceptualization: BS, LB, MT Methodology: JG, ASci ,ASch, FN Investigation: BS, FS, ASci, ASch, MF Funding acquisition: LB, FN, MT Supervision: ASci, ASch, LB, MT Writing – original draft: BS, FS, MT Writing – review & editing: all authors

## Competing interests

Authors declare that they have no competing interests.

---

**Figure S1:**
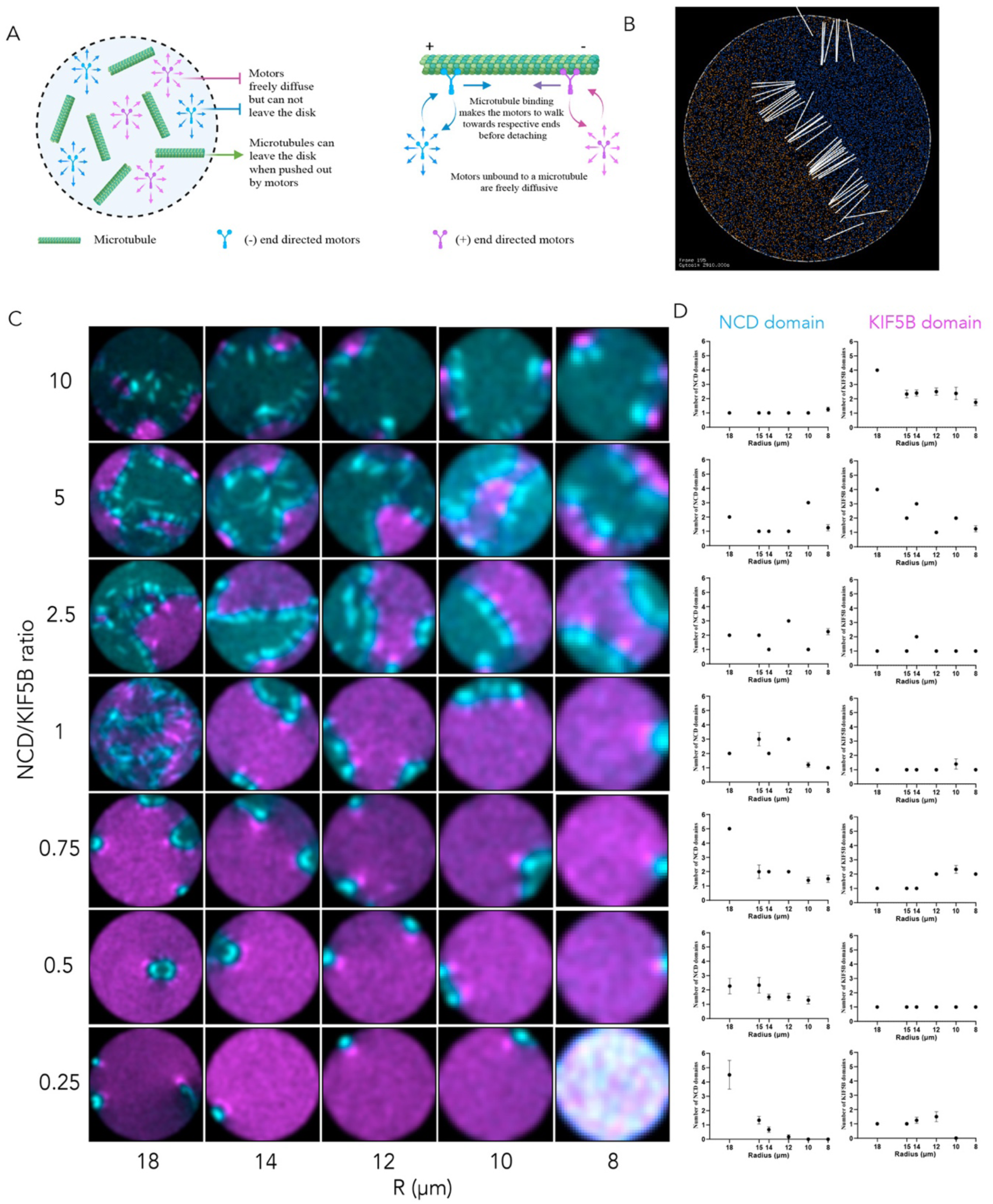
Agent-based simulations of microtubules and motors partitioning in confined area. A-Cartoon representation of the agent-based simulations under confinement. Motors were constrained to diffuse within a given area and could not escape from it. MTs were transported by the underlying motors but could be moved out of the confined region (see Materials and Methods). B-Example of simulation outcome with microtubules represented as white lines, and plus and minus-end motors as yellow and blue dots. C-Whole phase diagram (with motor fields-NCD is cyan and KIF5B is magenta) for motor domain formation by varying the relative motor ratio (y-axis) and confinement size (x-axis) using 2D modified-cytosim simulation (see Materials and Methods). X-axis represents the ratio of NCD to KIF5B and Y-axis defines the radius of the circle D-Quantification of the number of plus and minus-domains for the conditions tested in (C).

**Figure S2.**
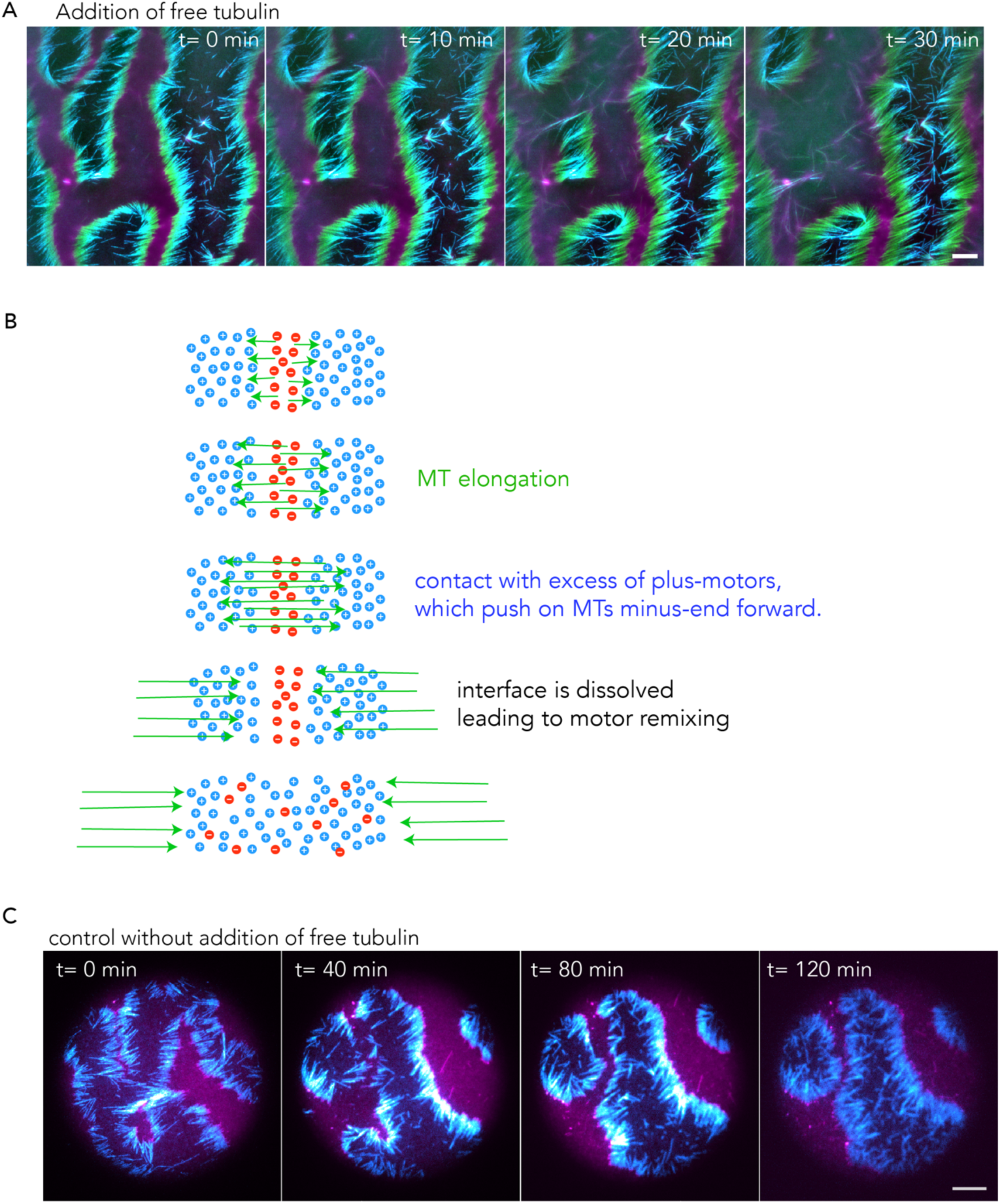
Microtubule elongation destabilizes the smallest pre-existing motor domains. A-Time-lapse fluorescence images showing microtubule elongation following the addition of free tubulin on an unconfined supported lipid bilayer (SLB). Images show microtubules (green), NCD (cyan), KIF5B (magenta), and their merged fluorescence signals. B-Schematic representation of how microtubule elongation disrupts the force balance between opposing motors, leading to the destabilization and dissolution of pre-existing motor domains. C-Control experiment showing time-lapse images of NCD (cyan) and KIF5B (magenta) domains within circular confinements in the absence of free tubulin addition. The motor domains remain organized over time without microtubule elongation. In all panels, scale bars represent 10 µm.

**Figure S3.**
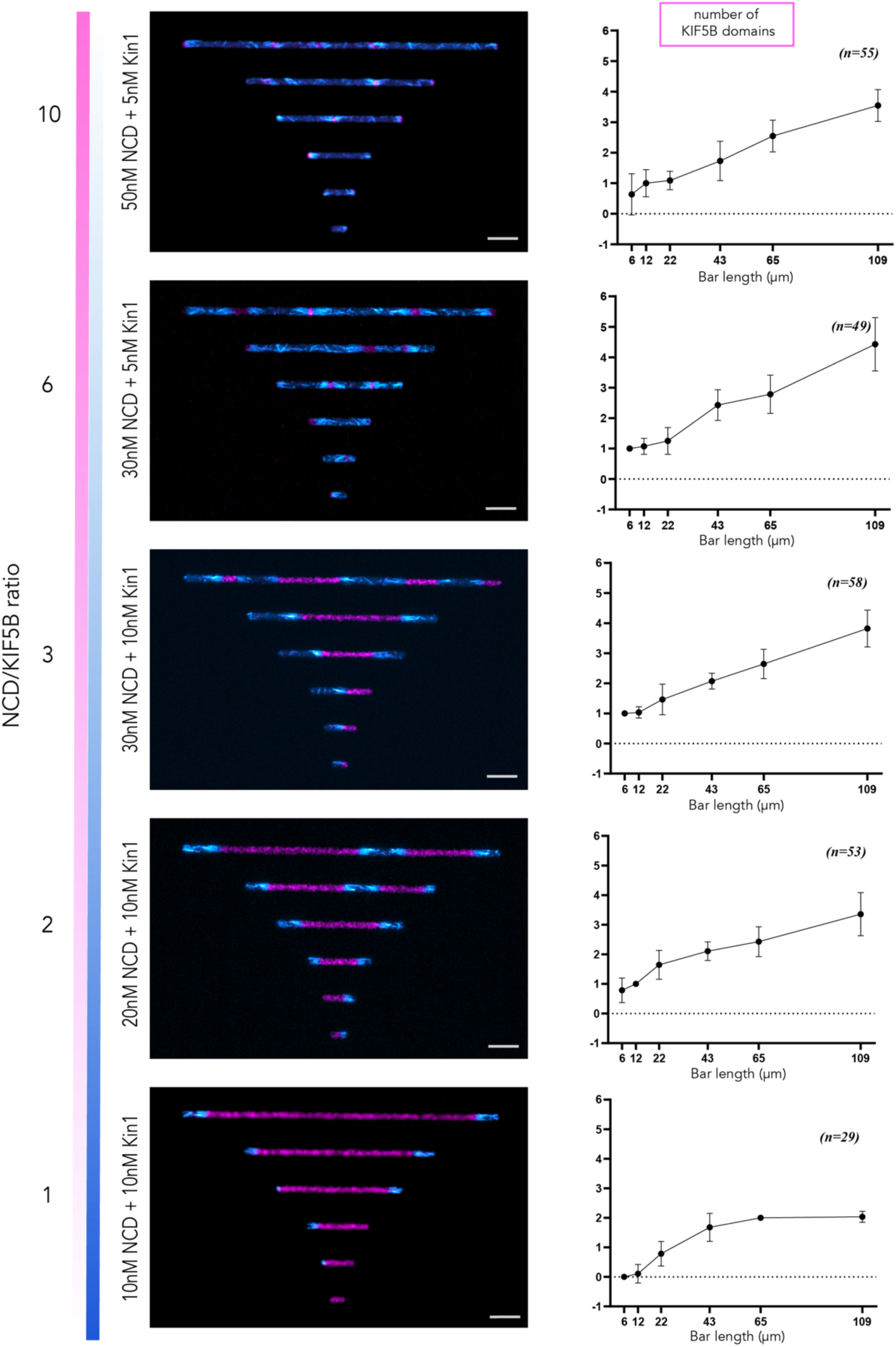
The number of partitions increases linearly with confinement length. Left: Fluorescence images showing the self-organization of KIF5B (magenta) and NCD (cyan) within elongated confinements of increasing length (6, 12, 22, 43, 65, and 109 µm) at varying NCD/KIF5B concentrations (10/10, 20/10, 30/10, 30/5, and 50/5 nM, bottom to top). Right: Quantification of the number of motor domains as a function of confinement length for the corresponding motor concentration combinations. (n = 29, 53, 58, 81, and 55 confinements per length for NCD/KIF5B concentrations of 10/10, 20/10, 30/10, 30/5, and 50/5 nM, respectively). Increasing confinement length allows the formation of additional motor domains across all motor ratios tested. Each point represents an individual confinement; horizontal lines and error bars represent the mean ± SD. Scale bars represent 10 µm.

**Figure S4.**
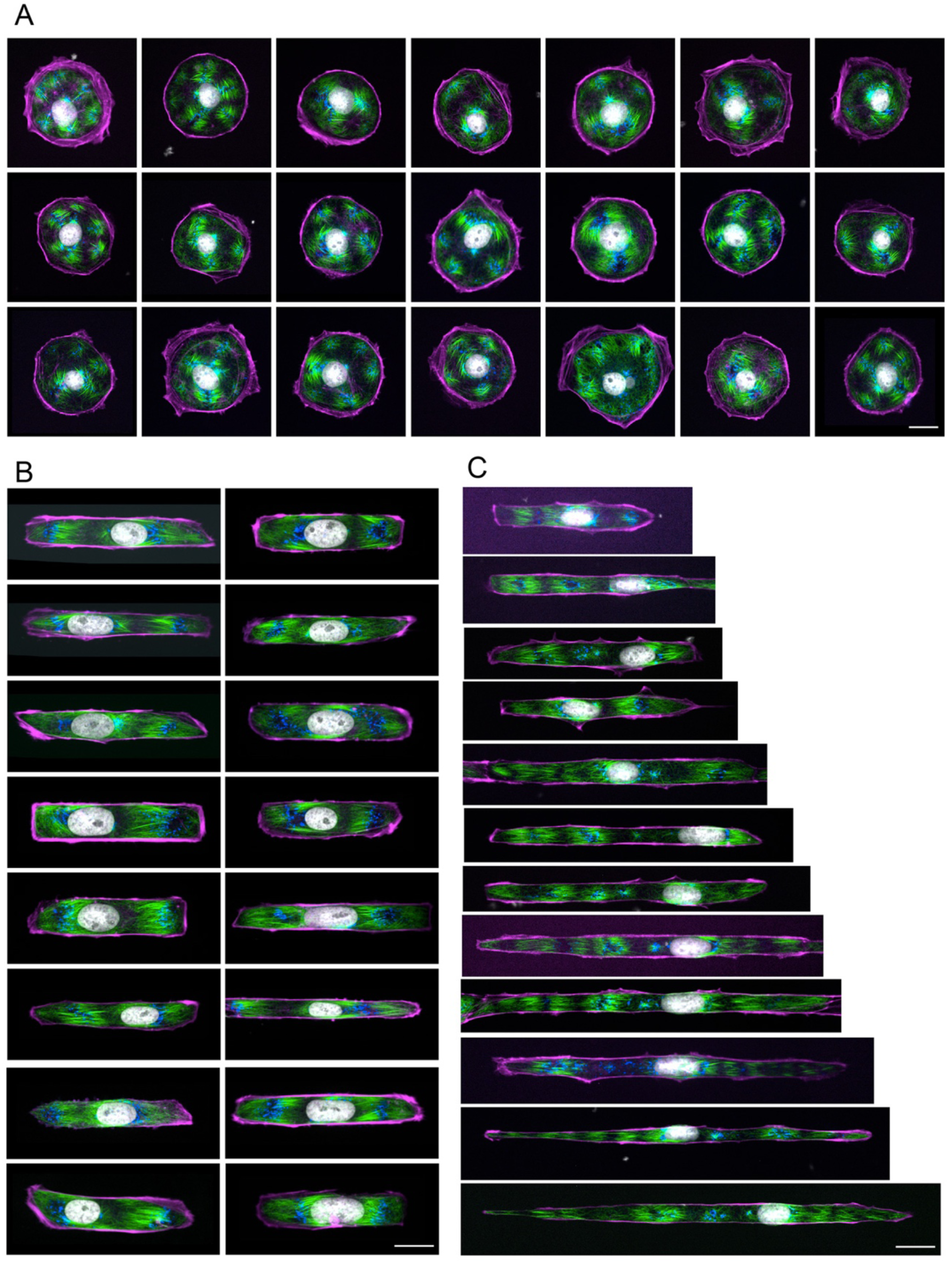
Spatial organization of partitions depends on cell shape aspect ratio. A,B,C-Immunofluorescence images showing actin (magenta), microtubules (green), Golgi apparatus (blue) and nucleus (gray) in fixed PtK2 cells. Cells were plated on 2000 µm² fibronectin-coated disc (A), or 12-µm-wide lines (B, C) micropatterns. Cells were then pretreated treated with 10µM Nocodazole 4 hours and then fixed 3h after Nocodazole washout and addition of 5µM Taxol. Cells were chosen to have either similar (B) or distinct length (C). In all panels, scale bars represent 20 µm.

## Notes

### Competing Interest Statement

The authors have declared no competing interest.

## References

1. G. J. Brouhard, L. M. Rice, Microtubule dynamics: an interplay of biochemistry and mechanics. Nat. Rev. Mol. Cell Biol., 1–13 (2018).

2. J. M. Cleary, W. O. Hancock, Molecular mechanisms underlying microtubule growth dynamics. Curr. Biol. 31, R560–R573 (2021).

3. A. Yildiz, Mechanism and regulation of kinesin motors. Nat. Rev. Mol. Cell Biol., doi: 10.1038/s41580-024-00780-6 (2024).

4. K. J. Verhey, J. W. Hammond, Traffic control: regulation of kinesin motors. Nat. Rev. Mol. Cell Biol. 10, 765–777 (2009).

5. S. D. Hunt, D. J. Stephens, The role of motor proteins in endosomal sorting. Biochem. Soc. Trans. 39, 1179–1184 (2011).

6. J. P. Caviston, E. L. F. Holzbaur, Microtubule motors at the intersection of trafficking and transport. Trends Cell Biol. 16, 530–537 (2006).

7. P. S. Gurel, A. L. Hatch, H. N. Higgs, Connecting the Cytoskeleton to the Endoplasmic Reticulum and Golgi. Curr. Biol. 24, R660–R672 (2014).

8. C. Gladkova, M. G. Paez-Segala, W. P. Grant, M. Kittisopikul, S. A. Myers, Y. Wang, R. D. Vale, A molecular switch for coordinating kinesin and dynein transport of mitochondrial cargo. Science 393, 601–606 (2026).

9. K. Barlan, V. I. Gelfand, Microtubule-Based Transport and the Distribution, Tethering, and Organization of Organelles. Cold Spring Harb. Perspect. Biol. 9, a025817–a025817 (2017).

10. J. Kroll, J. Renkawitz, Principles of organelle positioning in motile and non-motile cells. EMBO Rep. 25, 2172–2187 (2024).

11. M. Burute, L. C. Kapitein, Cellular logistics: Unraveling the interplay between microtubule organization and intracellular transport. Annu. Rev. Cell Dev. Biol. 35, 29–54 (2019).

12. A. Akhmanova, L. C. Kapitein, Mechanisms of microtubule organization in differentiated animal cells. Nat. Rev. Mol. Cell Biol. 23, 541–558 (2022).

13. C. E. Buckley, D. St Johnston, Apical–basal polarity and the control of epithelial form and function. Nat. Rev. Mol. Cell Biol. 23, 559–577 (2022).

14. A. N. Rao, P. W. Baas, Polarity Sorting of Microtubules in the Axon. Trends Neurosci. 41, 77–88 (2018).

15. M. Burute, M. Prioux, G. Blin, S. Truchet, G. Letort, Q. Tseng, T. Bessy, S. Lowell, J. Young, O. Filhol, M. Théry, Polarity Reversal by Centrosome Repositioning Primes Cell Scattering during Epithelial-to-Mesenchymal Transition. Dev. Cell 40, 168–184 (2017).

16. F. Peglion, S. Etienne-Manneville, Cell polarity changes in cancer initiation and progression. J. Cell Biol. 223, 1–15 (2024).

17. G. K. Ojakian, R. Schwimmer, K. G. Ojakian, R. Schwimmer, Regulation of epithelial cell surface polarity reversal by beta 1 integrins. J Cell Sci 107, 561–576 (1994).

18. M. K. Iwanski, A. K. Serweta, J. van Schelt, H. N. Verwei, B. C. Donders, L. C. Kapitein, Polarity reversal of stable microtubules during neuronal development. J. Cell Sci. 138 (2025).

19. J. Wu, A. Akhmanova, Microtubule-Organizing Centers. Annu. Rev. Cell Dev. Biol. 33, 51–75 (2017).

20. H. H. De Forges, A. Bouissou, F. Perez, Interplay between microtubule dynamics and intracellular organization. Int. J. Biochem. Cell Biol. 44, 266–274 (2012).

21. A. Akhmanova, M. O. Steinmetz, Control of microtubule organization and dynamics: two ends in the limelight. Nat. Rev. Mol. Cell Biol. 16, 711–726 (2015).

22. W. Lu, V. I. Gelfand, Moonlighting Motors: Kinesin, Dynein, and Cell Polarity. Trends Cell Biol. 27, 505–514 (2017).

23. C. E. Walczak, R. Heald, Mechanisms of mitotic spindle assembly and function. Int. Rev. Cytol. 265, 111– 58 (2008).

24. D. Oriola, D. J. Needleman, J. Brugués, The Physics of the Metaphase Spindle. Annu. Rev. Biophys. 47, 655–73 (2018).

25. A. L. Jolly, H. Kim, D. Srinivasan, M. Lakonishok, A. G. Larson, V. I. Gelfand, Kinesin-1 heavy chain mediates microtubule sliding to drive changes in cell shape. Proc. Natl. Acad. Sci. U. S. A. 107, 12151–6 (2010).

26. R. E. Carazo-Salas, C. Antony, P. Nurse, The Kinesin Klp2 Mediates Polarization of Interphase Microtubules in Fission Yeast. Science 309, 297–300 (2005).

27. R. E. Carazo-Salas, P. Nurse, Self-organization of interphase microtubule arrays in fission yeast. Nat. Cell Biol. 8, 1102–1107 (2006).

28. W. Lu, P. Fox, M. Lakonishok, M. W. Davidson, V. I. Gelfand, Initial neurite outgrowth in drosophila neurons is driven by kinesin-powered microtubule sliding. Curr. Biol. 23, 1018–1023 (2013).

29. W. Lu, M. Winding, M. Lakonishok, J. Wildonger, V. I. Gelfand, Microtubule–microtubule sliding by kinesin-1 is essential for normal cytoplasmic streaming in *Drosophila* oocytes. Proc. Natl. Acad. Sci. 113 (2016).

30. K. M. Bracey, M. A. Fye, A. Cario, K.-H. Ho, P. Noguchi, G. Gu, I. Kaverina, Glucose-stimulated KIF5B-driven microtubule sliding organizes microtubule networks in mouse pancreatic β cells. eLife 12, 1–25 (2025).

31. D. J. Needleman, J. Brugués, Determining Physical Principles of Subcellular Organization. Dev. Cell 29, 135–138 (2014).

32. L. Wordeman, How kinesin motor proteins drive mitotic spindle function: Lessons from molecular assays. Semin. Cell Dev. Biol. 21, 260–268 (2010).

33. D. J. Needleman, Z. Dogic, Active matter at the interface between materials science and cell biology. Nat. Rev. Mater. 2, 17048–17048 (2017).

34. F. Nedelec, T. Surrey, S. Leibler, A. C. Maggs, Self-organization of microtubules and motors. Nature 389, 305–308 (1997).

35. T. Surrey, F. Nedelec, S. Leibler, E. Karsenti, Physical properties determining self-organization of motors and microtubules. Science 292, 1167–71 (2001).

36. P. J. Foster, S. Fürthauer, M. J. Shelley, D. J. Needleman, S. Furthauer, M. J. Shelley, D. J. Needleman, Active contraction of microtubule networks. eLife 4, 1–21 (2015).

37. T. Sanchez, D. T. N. Chen, S. J. DeCamp, M. Heymann, Z. Dogic, Spontaneous motion in hierarchically assembled active matter. Nature, 1–5 (2012).

38. B. Najma, W.-S. Wei, A. Baskaran, P. J. Foster, G. Duclos, Microscopic interactions control a structural transition in active mixtures of microtubules and molecular motors. Proc. Natl. Acad. Sci. 121 (2024).

39. J. Roostalu, J. Rickman, C. Thomas, F. Nedelec, T. Surrey, Determinants of Polar versus Nematic Organization in Networks of Dynamic Microtubules and Mitotic Motors. Cell 175, 796–808.e14 (2018).

40. L. M. Lemma, M. M. Norton, A. M. Tayar, S. J. DeCamp, S. A. Aghvami, S. Fraden, M. F. Hagan, Z. Dogic, Multiscale Microtubule Dynamics in Active Nematics. Phys. Rev. Lett. 127, 1–13 (2021).

41. S. Fürthauer, B. Lemma, P. J. Foster, S. C. Ems-McClung, C. H. Yu, C. E. Walczak, Z. Dogic, D. J. Needleman, M. J. Shelley, Self-straining of actively crosslinked microtubule networks. Nat. Phys. 15, 1295–1300 (2019).

42. R. Jiang, S. Vandal, S. Park, S. Majd, E. Tüzel, W. O. Hancock, Microtubule binding kinetics of membrane-bound kinesin-1 predicts high motor copy numbers on intracellular cargo. Proc. Natl. Acad. Sci. U. S. A. 116, 26564–26570 (2019).

43. R. Grover, J. Fischer, F. W. Schwarz, W. J. Walter, P. Schwille, S. Diez, Transport efficiency of membrane-anchored kinesin-1 motors depends on motor density and diffusivity. Proc. Natl. Acad. Sci. U. S. A. 113, E7185–E7193 (2016).

44. D. Inoue, B. Mahmot, A. Md. R. Kabir, T. I. Farhana, K. Tokuraku, K. Sada, A. Konagaya, A. Kakugo, Depletion force induced collective motion of microtubules driven by kinesin. Nanoscale 7, 18054–18061 (2015).

45. F. L. Memarian, J. D. Lopes, F. J. Schwarzendahl, M. G. Athani, N. Sarpangala, A. Gopinathan, D. A. Beller, K. Dasbiswas, L. S. Hirst, Active nematic order and dynamic lane formation of microtubules driven by membrane-bound diffusing motors. Proc. Natl. Acad. Sci. U. S. A. 118, 1–10 (2021).

46. C. Hentrich, T. Surrey, Microtubule organization by the antagonistic mitotic motors kinesin-5 and kinesin-14. J. Cell Biol. 189, 465–80 (2010).

47. N. D. Derr, B. S. Goodman, R. Jungmann, A. E. Leschziner, W. M. Shih, S. L. Reck-Peterson, Tug-of-war in motor protein ensembles revealed with a programmable DNA origami scaffold. Science 338, 662–665 (2012).

48. J. Ikuta, N. K. Kamisetty, H. Shintaku, H. Kotera, T. Kon, R. Yokokawa, Tug-of-war of microtubule filaments at the boundary of a kinesin- and dynein-patterned surface. Sci. Rep. 4, 5281 (2014).

49. G. A. Monzon, L. Scharrel, A. DSouza, V. Henrichs, L. Santen, S. Diez, Stable tug-of-war between kinesin-1 and cytoplasmic dynein upon different ATP and roadblock concentrations. J. Cell Sci. 133, jcs249938– jcs249938 (2020).

50. A. I. D’Souza, R. Grover, G. A. Monzon, L. Santen, S. Diez, Vesicles driven by dynein and kinesin exhibit directional reversals without regulators. Nat. Commun. 14, 7532–7532 (2023).

51. C. Utzschneider, B. Suresh, A. Sciortino, J. Gaillard, A. Schaeffer, S. Pattanayak, J. F. Joanny, L. Blanchoin, M. Théry, Force balance of opposing diffusive motors generates polarity-sorted microtubule patterns. Proc. Natl. Acad. Sci. U. S. A. 121 (2024).

52. S. Pattanayak, A. Sciortino, L. Blanchoin, M. Théry, J.-F. Joanny, Phase Separation by Polar Active Transport. Phys. Rev. Lett. 137, 218302 (2026).

53. T. Vignaud, L. Blanchoin, M. Théry, Directed cytoskeleton self-organization. Trends Cell Biol. 22, 671–682 (2012).

54. M. Pinot, F. Chesnel, J. Z. J. Z. Kubiak, I. Arnal, F. Nedelec, Z. Gueroui, Effects of confinement on the self-organization of microtubules and motors. Curr. Biol. CB 19, 954–60 (2009).

55. M. P. N. Juniper, M. Weiss, I. Platzman, J. P. Spatz, T. Surrey, Spherical network contraction forms microtubule asters in confinement. Soft Matter 14, 901–909 (2018).

56. T. D. Ross, H. J. Lee, Z. Qu, R. A. Banks, R. Phillips, M. Thomson, Controlling organization and forces in active matter through optically defined boundaries. Nature 572, 224–229 (2019).

57. B. Najma, S. Ghosh, C. Amey, P. J. Foster, M. F. Hagan, A. Baskaran, G. Duclos, Arrested coalescence, aging, and stability of asters composed of microtubules and kinesin motors. Phys. Rev. Res. 7, 013247 (2025).

58. A. Opathalage, M. M. Norton, M. P. N. Juniper, B. Langeslay, S. A. Aghvami, S. Fraden, Z. Dogic, Self-organized dynamics and the transition to turbulence of confined active nematics. Proc. Natl. Acad. Sci. U. S. A. 116, 4788–4797 (2019).

59. K.-T. T. Wu, J. B. Hishamunda, D. T. N. N. Chen, S. J. DeCamp, Y.-W. W. Chang, A. Fernández-Nieves, S. Fraden, Z. Dogic, Transition from turbulent to coherent flows in confined three-dimensional active fluids. Science 355, eaal1979–eaal1979 (2017).

60. P. Chandrakar, M. Varghese, S. A. Aghvami, A. Baskaran, Z. Dogic, G. Duclos, Confinement Controls the Bend Instability of Three-Dimensional Active Liquid Crystals. Phys. Rev. Lett. 125, 257801–257801 (2020).

61. J. Hardoüin, J. Laurent, T. Lopez-Leon, J. Ignés-Mullol, F. Sagués, Active microfluidic transport in two-dimensional handlebodies. Soft Matter 16, 9230–9241 (2020).

62. A. Colin, M. Orhant-Prioux, C. Guérin, M. Savinov, W. Cao, B. Vianay, I. Scarfone, A. Roux, E. M. De La Cruz, A. Mogilner, M. Théry, L. Blanchoin, Friction patterns guide actin network contraction. Proc. Natl. Acad. Sci. 120, 2017–2017 (2023).

63. F. Verde, J.-M. Berrez, C. Antony, E. Karsenti, Taxol-induced Microtubule Asters in Mitotic Extracts of Xenopus Eggs : *J*. Cell Biol. 112, 1177–1187 (1991).

64. P. B. Schiff, S. B. Horwitz, Taxol stabilizes microtubules in mouse fibroblast cells. Proc. Natl. Acad. Sci. U. S. A. 77, 1561–1565 (1980).

65. K. J. Green, R. D. Goldman, The effects of taxol on cytoskeletal components in cultured fibroblasts and epithelial cells. Cell Motil. 3, 283–305 (1983).

66. G. Piperno, M. LeDizet, X. J. Chang, Microtubules containing acetylated alpha-tubulin in mammalian cells in culture. J. Cell Biol. 104, 289–302 (1987).

67. A. Harada, Y. Takei, Y. Kanai, Y. Tanaka, S. Nonaka, N. Hirokawa, Golgi vesiculation and lysosome dispersion in cells lacking cytoplasmic dynein. J. Cell Biol. 141, 51–59 (1998).

68. I. Corthésy-Theulaz, A. Pauloin, S. R. Pfeffer, I. Corthsy-theulaz, A. Pauloin, R. Pfeffer, Cytoplasmic dynein participates in the centrosomal localization of the Golgi complex. J. Cell Biol. 118, 1333–1345 (1992).

69. S. Yadav, M. A. A. Puthenveedu, A. D. D. Linstedt, Golgin160 Recruits the Dynein Motor to Position the Golgi Apparatus. Dev. Cell 23, 153–165 (2012).

70. C. Cadart, S. Monnier, J. Grilli, P. J. Sáez, N. Srivastava, R. Attia, E. Terriac, B. Baum, M. Cosentino-Lagomarsino, M. Piel, Size control in mammalian cells involves modulation of both growth rate and cell cycle duration. Nat. Commun. 9, 3275–3275 (2018).

71. J. Scrofani, F. Ruhnow, W.-X. Chew, D. Normanno, F. Nedelec, T. Surrey, I. Vernos, C. Walczak, Branched microtubule nucleation and dynein transport organize RanGTP asters in Xenopus laevis egg extract. Mol. Biol. Cell 35, ar12 (2024).

72. T. Fukuyama, L. Yan, M. Tanaka, M. Yamaoka, K. Saito, S.-C. Ti, C.-C. Liao, K.-C. Hsia, Y. T. Maeda, Y. Shimamoto, Morphological growth dynamics, mechanical stability, and active microtubule mechanics underlying spindle self-organization. Proc. Natl. Acad. Sci. 119, e2209053119 (2022).

73. F. Chen, J. Wu, M. K. Iwanski, D. Jurriens, A. Sandron, M. Pasolli, G. Puma, J. Z. Kromhout, C. Yang, W. Nijenhuis, L. C. Kapitein, F. Berger, A. Akhmanova, Self-assembly of pericentriolar material in interphase cells lacking centrioles. eLife 11 (2022).

74. S. R. Norris, S. Jung, P. Singh, C. E. Strothman, A. L. Erwin, M. D. Ohi, M. Zanic, R. Ohi, Microtubule minus- end aster organization is driven by processive HSET-tubulin clusters. Nat. Commun. 9, 1–14 (2018).

75. U. del Castillo, M. Winding, W. Lu, V. I. Gelfand, Interplay between kinesin-1 and cortical dynein during axonal outgrowth and microtubule organization in Drosophila neurons. eLife 4, 1–20 (2015).

76. D. B. Oelz, U. del Castillo, V. I. Gelfand, A. Mogilner, Microtubule Dynamics, Kinesin-1 Sliding, and Dynein Action Drive Growth of Cell Processes. Biophys. J. 115, 1614–1624 (2018).

77. M. Kwon, S. A. Godinho, N. S. Chandhok, N. J. Ganem, A. Azioune, M. Thery, D. Pellman, Mechanisms to suppress multipolar divisions in cancer cells with extra centrosomes. Genes Dev. 22, 2189–2203 (2008).

78. C. So, K. Menelaou, J. Uraji, K. Harasimov, A. M. Steyer, K. B. Seres, J. Bucevičius, G. Lukinavičius, W. Möbius, C. Sibold, A. Tandler-Schneider, H. Eckel, R. Moltrecht, M. Blayney, K. Elder, M. Schuh, Mechanism of spindle pole organization and instability in human oocytes. Science 375 (2022).

79. D. Clift, M. Schuh, A three-step MTOC fragmentation mechanism facilitates bipolar spindle assembly in mouse oocytes. Nat. Commun. 6, 1–12 (2015).

80. H. Musa, C. Orton, E. E. Morrison, M. Peckham, Microtubule assembly in cultured myoblasts and myotubes following nocodazole induced microtubule depolymerisation. J. Muscle Res. Cell Motil. 24, 301–308 (2003).

81. A. M. Tassin, B. Maro, M. Bornens, Fate of microtubule-organizing centers during myogenesis in vitro. J. Cell Biol. 100, 35–46 (1985).

82. B. Cadot, V. Gache, E. Vasyutina, S. Falcone, C. Birchmeier, E. R. Gomes, Nuclear movement during myotube formation is microtubule and dynein dependent and is regulated by Cdc42, Par6 and Par3. EMBO Rep. 13, 741–749 (2012).

83. T. Metzger, V. Gache, M. Xu, B. Cadot, E. S. Folker, B. E. Richardson, E. R. Gomes, M. K. Baylies, MAP and kinesin-dependent nuclear positioning is required for skeletal muscle function. Nature 484, 120–124 (2012).

84. V. Gache, E. R. Gomes, B. Cadot, Microtubule motors involved in nuclear movement during skeletal muscle differentiation. Mol. Biol. Cell 28, 865–874 (2017).

85. M. H. Wilson, E. L. F. Holzbaur, Nesprins anchor kinesin-1 motors to the nucleus to drive nuclear distribution in muscle cells. Development 142, 218–228 (2015).

86. H. Liu, J. P. I. Welburn, A circle of life : platelet and megakaryocyte cytoskeleton dynamics in health and disease. (2024).

87. R. Philip, C. Fiorino, R. E. Harrison, Terminally differentiated osteoclasts organize centrosomes into large clusters for microtubule nucleation and bone resorption. Mol. Biol. Cell 33, ar68 (2022).

88. C. A. Anderson, U. Eser, T. Korndorf, M. E. Borsuk, J. M. Skotheim, A. S. Gladfelter, Nuclear repulsion enables division autonomy in a single cytoplasm. Curr. Biol. 23, 1999–2010 (2013).

89. S. Grava, M. Keller, S. Voegeli, S. Seger, C. Lang, P. Philippsen, Clustering of Nuclei in Multinucleated Hyphae Is Prevented by Dynein-Driven Bidirectional Nuclear Movements and Microtubule Growth Control in Ashbya gossypii ▿. Eukaryot. Cell 10, 902–915 (2011).

90. J. De-Carvalho, S. Tlili, T. E. Saunders, I. A. Telley, The positioning mechanics of microtubule asters in Drosophila embryo explants. eLife 12, 1–33 (2024).

91. X. Cheng, J. E. Ferrell, Spontaneous emergence of cell-like organization in Xenopus egg extracts. Science 366, 631–637 (2019).

92. D. Nashchekin, L. Busby, M. Jakobs, I. Squires, D. St. Johnston, Symmetry breaking in the female germline cyst. Science 374, 874–879 (2021).

93. W. Lu, M. Lakonishok, V. I. Gelfand, The dynamic duo of microtubule polymerase Mini spindles/XMAP215 and cytoplasmic dynein is essential for maintaining *Drosophila* oocyte fate. Proc. Natl. Acad. Sci. 120, e2303376120 (2023).

