## Supplementary Information for "Space partitioning by microtubules and molecular motors"

- Materials and Methods
- Supplementary References
- Supplementary movies

### Materials and Methods

#### In vitro experiments with microtubules and motors

##### Protein expression, purification and labelling

**Tubulin**

Bovine brain tubulin was purified in BRB80 buffer (80 mM 1,4-piperazinediethanesulfonic acid, pH 6.8, 1 mM ethylene glycol tetra acetic acid, and 1 mM MgCl2) according to previously published method (*1*). Tubulin was purified from fresh bovine brain by three cycles of temperature-dependent assembly and disassembly in BRB80 supplemented with 1 mM GTP (*2*). MAP-free neuro-tubulin was purified using cation-exchange chromatography (EMD SO, 650 M, Merck) in 50 mM PIPES, pH 6.8, supplemented with 1 mM MgCl2, and 1 mM EGTA. Purified tubulin was obtained after a cycle of polymerization and depolymerization. Fluorescent tubulin (ATTO 488–labeled tubulin and ATTO 565–labeled tubulin) and biotinylated tubulin were prepared according to previously published method (*3*). Microtubules from neuro-tubulin were polymerized at 37°C for 30 min and layered onto cushions of 0.1 M Na-HEPES, pH 8.6, 1 mM MgCl2, 1 mM EGTA, 60% v/v glycerol, and sedimented by high centrifugation at 30°C. Then microtubules were resuspended in 0.1 M Na-HEPES pH 8.6, 1 mM MgCl2, 1 mM EGTA, 40% v/v glycerol and labeled by adding 1/10 volume 100 mM NHS-ATTO (ATTO Tec) for 10 min at 37°C. The labeling reaction was stopped using 2 volumes of 2X BRB80, containing 100 mM potassium glutamate and 40% v/v glycerol, and then microtubules were sedimented onto cushions of BRB80 supplemented with 60% glycerol. Microtubules were resuspended in cold BRB80. Microtubules were then depolymerized and a second cycle of polymerization and depolymerization was performed. Tubulin was kept in liquid nitrogen. Our labelling ratio is: for tubulin 488 nm 1.6 to 1.8 fluorophore per dimer, and for tubulin 565 nm 0.6 to 0.7 fluorophore per dimer. We used NanoDrop one UV-Vis spectrophotometer (Thermofisher) to determine the labelling efficiency.

**NCD**

Recombinant, truncated NCD motor protein (amino acid 236-700, accession number A0A0B4LI25) was purified using bacterial expression as previously described (*4*). The construct includes sequences encoding a His6 tag and GFP at the N-terminal end of the protein sequence. Expression of the plasmid, pET17b_NCD_GFP_His (24507, Addgene, Cambridge, MA) was induced by transfection into Rosetta2 (DE3)-pLysS *E. coli* (VWR), followed by induction with 0.2 mM IPTG for 16 h at 18°C.

The purification of the expressed kinesin was performed in three steps. First, the raw lysate was loaded onto a Ni-NTA column to affinity purify via the N-terminal histidine tag. The flow-through from this column was then slightly diluted and loaded onto an anion exchange column. The final purification step involved gel filtration. The motors were stored frozen at -80°C.

Procedure for Ni-NTA column purification: The pellet was lysed in 80 ml of lysis buffer (50 mM NaPO4, pH 8.0, 20 mM imidazole, 250 mM NaCl, 1 mM MgCl2, 0.5 mM ATP, 10 mM β-mercaproethanol, 1x protease inhibitor cocktail (complete, EDTA free, Roche, Mannheim, Germany)) and centrifuged at 25000g for 30 min at 4°C. The supernatant (lysate) was loaded onto a Ni-NTA column (HisTrap HP$^{TM}$, 17-5247-01, Cytiva), washed with washing buffer (50 mM NaPO4, pH 6.0, 250 mM NaCl, 1 mM MgCl2, 0.1 mM ATP, 10 mM β-

mercaproethanol), and eluted with elution buffer (50 mM NaPO4, pH 7.2, 500 mM Imidazole, 250 mM NaCl, 1 mM MgCl2, 0.1 mM ATP, 10 mM β-mercaproethanol).
Procedure for the gel filtration column: The eluate from the anion exchange column was loaded onto a gel filtration column (hiload 16 600 superdex 200 pg, 28-9893-35, Cytiva) and eluted with NCD buffer (10 mM NaPO4, pH 7.2, 50 mM NaCl, 2 mM MgCl2, 1 mM EGTA, 1 mM DTT, 0.1 mM ATP).

**KIF5B**

Recombinant, truncated kinesin-1 motor protein (amino acids 1–560, accession number : P33176 ) was purified using bacterial expression as previously described (*5*). The construct includes the sequences encoding a His6 tag and SNAP tag at the C-terminal end of the protein sequence. The plasmid for the kinesin construct, pET17_K560_SNAP_His, was generously provided by Dr. Ross. The plasmid was introduced into Rosetta2 (DE3)-pLysS *E. coli* (VWR) by transformation, and induction was carried out with 0.2 mM IPTG for 16 hours at 18°C.

The purification of the expressed kinesin was performed in three steps. The first step involves using a cation exchange column. The bacteria pellet was lysed in 80 ml lysis buffer (Lysis buffer is a cation exchange buffer (6.7 mM sodium acetate, 6.7 mM 4-(2-hydroxyethyl)-1-piperazineethanesulfonic acid (HEPES), 6.7 mM 2-ethanesulfonic acid (MES), pH 7.0, 20 mM beta mercaptoethanol (BME), 0.2 mM ATP, 0.2% (w/v) polyoxyethylene (20) sorbitan monolaurate (TWEEN20)) supplemented with 1x protease inhibitor cocktail (complete, EDTA free, Roche, Mannheim, Germany) and TWEEN20 to a final concentration of 0.5% (w/v)). After sonication, and centrifugation at 25,000 g for 30 minutes at 4°C, the supernatant (lysate) was loaded onto a cation exchange column (HiTrap SP HP™, 17-1151-01, Cytiva), washed with washing buffer (cation exchange buffer supplemented with KCl to a final concentration of 50 mM) and eluted with elution buffer (cation exchange buffer supplemented with KCl to a final concentration of 300 mM).

The flow-through from that column was diluted 5-fold in nickel loading buffer (nickel buffer: 50 mM sodium phosphate buffer, pH 7.5, 5% w/v glycerol, 300 mM KCl, 1 mM MgCl2, 0.2% w/v TWEEN20, 10 mM BME, 0.1 mM ATP, supplemented with imidazole to a final concentration of 36 mM) and loaded onto a Ni-NTA column (HisTrap HP™, 17-5247-01, Cytiva), washed with nickel washing buffer (nickel buffer supplemented with KCl to a final concentration of 1000 mM and imidazole to a final concentration of 30 mM) and eluted with nickel elution buffer (nickel buffer supplemented with imidazole to a final concentration of 300 mM). We added SNAP surface Alexa Fluor 647 (S9136,New England Biolabs) at the same concentration as kinesin. The SNAP tag was labeled overnight at 4°C. Fluorescently-labeled kinesin was loaded onto a gel filtration column (Superdex 200 Increase 10/300 GL, 28-9909-44, Cytiva) and eluted with K560 buffer (50 mM NaPO4 (pH7.4), 300 mM KCl, 1 mM MgCl2, 5% glycerol, 1mM DTT, 0.1mM ATP).

#### Lipids and SUV preparation

First, 96% L-α-phosphatidylcholine (Egg-PC) (Avanti, 840051C) (10 mg/mL in chloroform), 2% DGS-NTA(Ni) (Avanti, 790404C), and 1% DOPE-ATTO390 (ATTO-TEC, 390-161) were mixed into glass tubes, dried with argon gas, and incubated under vacuum overnight. The dried lipids were then resuspended in a buffer that we further refer to as the "small unilamellar vesicle (SUV) buffer" (150 mM NaCl, 10 mM Tris-HCl pH 7.4 and 2 mM CaCl2) and sonicated on ice for 10 min to generate the SUV. The mixture was centrifuged at

20,238 g for 10 min to remove contaminants and the supernatant was collected. The final lipid concentration was 1 mg/ml.

#### Stabilized microtubules preparation

Purified tubulin (20 µM) were mixed in BRB80 and incubated at 37 °C for 2 h. For the preparation of labelled microtubules, the labelled tubulin was added, with a final concentration of 20 µM into the polymerization mixture, The mixture was then centrifuged at 20,238×g for 30 min and the microtubules were resuspended in BRB80 supplemented with 10 µM of Taxol. Stabilized microtubules were kept at room temperature for 4 days.

#### Surface passivation and micropatterning

Glass coverslips were cleaned by sonication in 2% Hellmanex III for 30 min at 60 °C, rinsed with ultrapure water, dried with compressed air, and activated with air plasma for 5 min. 250 ml of 96% ethanol was taken in a clean glass beaker and stirred with a magnetic bead at 69 °C. Then, 670 µl of 37% HCl was added with a glass dropper to make it into 0.1% final solution. Keep the silane PEG 30K at room temperature for few minutes before opening to avoid the humidity. 0.25 g of silane PEG 30K was measured and then added to the Ethanol-HCl solution. It was covered with an aluminum foil and stirred for 1 hour until the PEG is completely dissolved. This solution can be kept at room temperature for a month. The coverslips were then incubated overnight in this solution of silane-PEG (30 kDa) to passivate the glass surface. Following silanization, the coverslips were rinsed with 96% ethanol, washed twice with ultrapure water, and dried with compressed air.

When necessary, PDMS wells were used to allow protein addition during the experiment without generating the flow associated to flow chambers. They were prepared by punching a 5-mm-diameter hole into cured PDMS and cutting a square around the opening. The PDMS wells were activated with air plasma for 30 s and gently pressed onto the silane-PEG-coated coverslips.

Micropatterns of defined sizes and geometries were primarily generated using maskless photolithography with the PRIMO micropatterning system (Alvéole). The desired micropattern geometries were designed using Inkscape and exported as PNG files. The PEGlated glass slides were covered with 20 µL of PLPP, a benzophenone-based photoinitiator reagent (Alvéole). Photopatterning was performed using a Nikon Plan Fluor 20× objective, with a calibrated pixel size of 0.286 µm/pixel and a UV exposure dose of 100 mJ/mm². Following UV exposure, the PLPP solution was removed, and the slides were rinsed five times with 20 µL of small unilamellar vesicle (SUV) buffer. Alternatively, micropatterns were generated by deep-UV exposure through a chrome photomask. The cleaned photomask was placed in contact with the silane-PEG-coated coverslip, held in position under vacuum, and exposed to UV light for 30 s.

In both cases, supported lipid bilayers were formed on the micropatterned surfaces by adding the SUV solution described above. In contact with micropatterned regions, SUVs spread, rupture and cover them with their lipid bilayer.

#### Motor-domain formation experiments

Cover glasses and slides were first wiped with lint-free tissues and Ethanol 96%, rinsed in ultrapure water, put in Hellmanex III (2% in water, Hellmanex), and sonicated for 30 min at 60 °C. The glass was then rinsed in ultrapure water and kept in water for 4 days. We have used open PDMS microwells to perform most experiments with occassional use of flow cells. In both cases, the bottom glass coverslip was dried with compressed air, activated with air plasma for 3 min. In the case of open microwells, a PDMS square with a punched out 5 mm hole in the center was placed on the glass creating a well. In the case of flow cells, two bands of double-side tape (70 µm height) were covered by a glass coverslip to contain an approximate volume of 15 µl. In both cases, 20 µL of lipids was added, incubated for 10 min then rinsed with SUV buffer. For additional passivation, 1% w/v BSA and 0.5 mg/ml PLL-PEG in HKEM (10 mM HEPES pH = 7.2, 50 mM KCl, 1 mM EGTA, 5 mM MgCl2) was incubated for 5 min. BSA was washed out with Tic-tac buffer (10 mM HEPES buffer (pH 7.2), 16 mM PIPES buffer (pH 6.8), 50 mM KCl, 5 mM MgCl2, 1 mM EGTA, 20 mM dithiothreitol (DTT), 3 mg.ml–1 glucose, 20 µg.ml–1 catalase, 100 µg.mL–1 glucose oxidase, 0.3% BSA) supplemented with 1 mM ATP (Sigma Aldrich, A3377). Motors diluted in tic-tac buffer were added and incubated for 5 min, then washed. Finally, tic-tac buffer supplemented with 2.7 mM of ATP and microtubules were added prior to imaging. Experiments were performed at room temperature.

### In vivo experiments with cells

**Cell culture**

PtK2 (Potorous tridactylus kidney) cells (control or stably expressing GFP-alpha-tubulin) (Khodjakov et al., 2003) were obtained from the lab of Franck Perez and were not further characterized. Cells were grown at 37°C and 5% CO2 in DMEM/F12 supplemented with 10% fetal bovine serum and 1% of Antibiotic-Antimycotic solution (ThermoFisher). Upon passage, cells were detached using TrypLE (ThermoFisher).

**Coverslip micropatterning**

Polystyrene coating

Glass slides were prepared for cell adhesion by coating them with polystyrene. Prior to coating, the slides were cleaned and treated with air plasma for 3 min (PE-50, Plasmatech). They were then exposed to HMDS vapor in a vacuum chamber for 1 h. Following this treatment, a 0,5% polystyrene solution in toluene was deposited onto the slides using a spin coater (WS-650m2-23NPPB, Laurell). The coating was performed at 1,500 rpm for 30 s. The coated slides were left to dry for at least 1 h before use and could be stored for several weeks in a dry, light-protected environment. Slides were then micropatterned either with deep-UV or normal UV and a photo-initiator

Deep-UV exposure

Polystyrene coated slides were activated using a 45 second air plasma treatment (PE50 plasmatech). After activation, the slides were then placed inside a solution of PLLPEG for at least 30 minutes at room temperature. The coating solution consisted of 0.1 mg/ml poly(L-lysine)-poly(ethylene glycol) (PLL-PEG, jenkem Technology) prepared in 10 mm HEPES buffer (ph 7.4). Following incubation, the slides were briefly rinsed in Milli-Q water and immediately dried. The coated slides were then stored at 4°C for at least 1 h before use.

PLL-PEG-coated coverslips were covered with a quartz–chrome photomask (Toppan Photomask) and held in close contact using a vacuum holder. Deep UV exposure (180 nm) was performed through the mask's transparent regions using a UVO cleaner (Model 342A-220, Jelight). The samples were positioned 1 cm from the UV source and irradiated at an intensity of 6 mw/cm² for 5 min. Following exposure, the coverslips were released from the photomask by immersion in Milli-Q water. The samples were then detached, dried, and incubated in an adhesion-protein solution according to the protein-coating procedure.

Normal UV exposure

Polystyrene coated coverslips were first coated with poly-L-lysine (PLL, 0.01%, Sigma Aldrich P4832) 30 minutes at room temperature, and then passivated with 100 mg/mL mPEG-SVA (5kDa, Laysan Bio) in 100 mM HEPES buffer (pH 8). PLPP-gel (Alvéole) was then deposited onto the surface as a photoinitiator, and the samples were transferred to an inverted microscope (Ti-E, Nikon) equipped with the PRIMO system (Alvéole). Stage-to-camera alignment and calibration of the Digital Micromirror Device (DMD) were performed using Leonardo software (Alvéole). UV illumination was applied at a dose of 30 mj/mm² to locally remove the passivation coating and generate the desired micropatterns. Finally, the micropatterned slides were incubated with fibronectin (10 µg/ml) for 20 min at room temperature to promote cell adhesion.

**Fixation and immunostaining**

Cells were fixed in methanol for 3 min at −20°C and subsequently washed three times with PBS. The samples were then incubated in a blocking solution (PBS, 0.1% Tween-20, 3% BSA (A2153, Sigma)) for 30 min at RT. Primary antibodies directed against tyrosinated tubulin (YL1/2: MAB1864, Merck at 1 :300), GM130 (MA5-35107 at 1 :200), β-actin (66009, Proteintech at 1 :400) were diluted in blocking solution and incubated with the samples for 30 min at RT. The slides were then rinsed three times with PBS-Tween 0.1% and incubated for 30 min at RT with the appropriate secondary antibodies (711-605-152, Jackson immunoresearch, anti-rabbit at 1:300; 712-545-143, Jackson immunoresearch, anti-rat at 1 :300; A31570, Invitrogen, anti-mouse at 1 :300), in blocking solution; DAPI (D9542, Sigma) was added at a 1/2000 dilution. Finally, the slides were rinsed twice with PBS-Tween 0.1%, once with water, and mounted in Mowiol (10981, Sigma).

#### Microscopy for microtubule networks and cells

To image networks of microtubule and motors, fluorescence imaging was performed using two microscopy platforms depending on the experimental requirements. Widefield fluorescence imaging was performed on a Nikon Eclipse Ti2 microscope equipped with a 60× oil objective, a Prime BSI-Express camera (pixel size = 6.5 µm, binning = 2), and 488, 561, and 642 nm lasers. Total internal reflection fluorescence (TIRF) imaging was carried out either on inverted Nikon Ti-E microscope equipped with an iLasPulsed TIRF system (Gataca Systems), a Retiga R3 CCD camera (1920 × 1460 pixels, binning = 1, pixel size = 4.54 µm), and an Olympus U APO N TIRF oil-immersion 100×/1.49 NA objective fitted with a 0.63 O-ring. Multi-stage time-lapse recordings were acquired using MetaMorph software (version 7.7.5, Universal Imaging) with a general exposure time of 100 ms per channel, at 10% laser power and variable time of acquisitions depending on the experiments.

To image cells, acquisitions were made with a confocal spinning disk microscope (Nikon Ti Eclipse equipped with a spinning scanning unit CSU-X1 Yokogawa) and a R3 retiga camera (qimaging). Images were acquired using either Nikon CFI Plan Fluor 40x/1.3 NA oil objective, Nikon Plan Apo VC 60x/1.40 NA oil objective, objective. Each wavelength was acquired separately. Metamorph software was used for images acquisition. Live acquisitions were performed inside a live module (kept at 37°C and 5% $CO_2$) mounted on the confocal spinning disk microscope.

### In silico experiments

#### Agent based simulations

We simulated microtubules interacting with two populations of oppositely directed molecular motors using Cytosim, an open-source framework for stochastic simulations of cytoskeletal mechanics (*6*, *7*). The two-dimensional model represents individual filaments and motors, motivated by microtubule gliding assays with motors bound to a fluid supported lipid bilayer (*8*, *9*). The membrane is implicit; its geometry and motor mobility are specified through the pattern boundary and motor parameters. The source code has been modified to allow motors to diffuse also while bound by integrating the force-balanced equation of motion of a motor bound to a filament even if the cytosim parameter activity is set to "fixed" (immobile motor) (*10*).

Microtubules were represented as polar, semiflexible, inextensible fibers with a bending rigidity of 10 pN $\mu m^2$, discretized with a target segment length of 0.1 µm. Translation, rotation, and bending resulted from thermal fluctuations, viscous drag, motor forces, and short-range steric repulsion. Cytosim integrates the constrained overdamped Langevin equations using an implicit treatment of mechanical interactions [1]. Filament motion was therefore calculated dynamically rather than imposed as a prescribed gliding velocity. The effective viscosity was 20 pN s $\mu m^{-2}$ and the thermal energy was 0.00411 pN µm.

Each motor was represented by a single object carrying one microtubule-binding hand. The two species were plus-end-directed kinesin and minus-end-directed Ncd. Both were assigned a diffusion coefficient of 0.75 $\mu m^2\ s^{-1}$. A motor within 0.05 µm of a filament could bind stochastically with a rate of 5 $s^{-1}$; this is an encounter-conditioned rate, not a bulk bimolecular association constant. Attachment created a Hookean link of stiffness 30 pN $\mu m^{-1}$ between the motor anchor and its filament-binding site. Bound heads moved continuously along the filament with a linear force–velocity relation, while the resulting spring forces displaced and deformed the microtubule. Attached-anchor mobility is qualified in the implementation notes.

The signed unloaded head velocities were +1.0 $\mu m\ s^{-1}$ for kinesin and −0.2 $\mu m\ s^{-1}$ for Ncd, with positive motion directed toward the microtubule plus end. Both species had a stall force of 6 pN. Detachment was force-dependent, where an unbinding force of 10 pN and a force-free off rate of 0.1 $s^{-1}$ for kinesin and 0.75 $s^{-1}$ for Ncd. Each motor had one filament-binding hand, so motor-mediated crosslinking of two microtubules was not specified. Growth-end tracking was disabled for both motor species.

Microtubules were initialized as 0.05-µm seeds inside the pattern. Their plus ends elongated at an unloaded rate of 0.10 $\mu m\ s^{-1}$ up to a maximum length of 3.5 µm; minus-end growth and shortening at both ends were set to zero. Filaments evolved for 45 s to reach the final length before both motor populations were introduced. Motor–microtubule dynamics were then simulated for 3,600 s, using a time step of 0.001 s throughout. Motors were initially placed inside the pattern. No mechanical confinement was specified for microtubules: the pattern boundary exerted no explicit filament-wall restoring force so filaments can live the pattern if pushed out. An event with delay 45 s removed all filaments outside the pattern. Motors are instead confined in the pattern.

Motor and microtubule numbers, pattern radius, and pattern shape were varied between conditions.

### Simulation and motor parameters

| Simulation parameter | Value |
|---|---|
| Spatial dimension | 2 |
| Time step | 0.001 s |
| Effective viscosity | 20 pN s $\mu m^{-2}$ |
| Thermal energy | 0.00411 pN µm |
| Pattern shape | **varying** |
| Pattern radius or characteristic size | **varying** |
| Global steric setting | 1, 400, 0 |
| Maximum steric search range | 0.1 µm |
| Initialization before motor addition | 45 s / 45,000 steps |
| Dynamics after motor addition | 3,600 s / 3,600,000 steps |
| Saved frames in initialization and main run | 3 and 240 |
| Fiber-removal event delay | 45 s |

#### Motor parameters

| Parameter | Motor 1 kinesin | Motor 2 Ncd |
|---|---|---|
| Number | **varying** | **varying** |
| Direction | Plus end | Minus end |
| Representation | single with one hand | single with one hand |
| Hand activity | move | move |
| Single activity[1] | fixed | fixed |
| Specified diffusion coefficient[1] | 0.75 $\mu m^2\ s^{-1}$ | 0.75 $\mu m^2\ s^{-1}$ |
| Binding rate | 5 $s^{-1}$ | 5 $s^{-1}$ |
| Binding range | 0.05 µm | 0.05 µm |
| Zero-load detachment rate | 0.1 $s^{-1}$ | 0.75 $s^{-1}$ |
| Detachment force scale | 10 pN | 10 pN |
| Signed unloaded velocity | +1.0 µm $s^{-1}$ | −0.2 µm $s^{-1}$ |
| Stall force | 6 pN | 6 pN |
| Link stiffness | 30 pN $\mu m^{-1}$ | 30 pN $\mu m^{-1}$ |
| Hold growing end | 0 | 0 |
| Initial placement | inside | inside |
| Confinement setting | inside, 0, cell | inside, 0, cell |

### Microtubule parameters

| Parameter | Value |
|---|---|
| Initial number | **varying** |
| Bending rigidity | 10 pN $\mu m^2$ |
| Target segment length | 0.1 µm |
| Initial seed length | 0.05 µm |
| Maximum length | 3.5 µm |
| Length-dynamics activity | grow |
| Initial end states | green, white |
| Plus- and minus-end growth speeds | 0.10, 0 $\mu m\ s^{-1}$ |
| Plus- and minus-end shortening speeds | 0, 0 $\mu m\ s^{-1}$ |
| Plus- and minus-end growth force scales | 3, 3 pN |
| Fiber steric setting[2] | 1, 0.0200, 0.0400 |
| Mechanical boundary confinement | Not specified |
| Boundary-related removal | Selection by position = outside, cell |

### Supplementary References


1. M. Vantard, C. Peter, A. Fellous, P. Schellenbaum, A. M. Lambert, Characterization of a 100-kDa heat-stable microtubule-associated protein from higher plants. *Fed. Eur. Biochem. Soc. J.* **220**, 847–853 (1994).
2. M. L. Shelanski, Chemistry of the filaments and tubules of brain. *J. Histochem. Cytochem. Off. J. Histochem. Soc.* **21**, 529–39 (1973).
3. A. Hyman, D. Drechsel, D. Kellogg, S. Salser, K. Sawin, P. Steffen, L. Wordeman, T. J. Mitchison, Preparation of modified tubulins. *Methods Enzymol.* **196**, 478–485 (1991).
4. R. B. Case, D. W. Pierce, N. Hom-Booher, C. L. Hart, R. D. Vale, The directional preference of kinesin motors is specified by an element outside of the motor catalytic domain. *Cell* **90**, 959–966 (1997).
5. T. Korten, S. Chaudhuri, E. Tavkin, M. Braun, S. Diez, Kinesin-1 Expressed in Insect Cells Improves Microtubule in Vitro Gliding Performance, Long-Term Stability and Guiding Efficiency in Nanostructures. *IEEE Trans. Nanobioscience* **15**, 62–69 (2016).
6. F. Nedelec, D. Foethke, Collective Langevin dynamics of flexible cytoskeletal fibers. *New J. Phys.* **9**, 427–427 (2007).
7. EMBL, Simulating the Cytoskeleton Using Cytosim. https://embl.org/training/self-paced-learning/simulating-cytoskeleton-cytosim/.
8. C. Utzschneider, B. Suresh, A. Sciortino, J. Gaillard, A. Schaeffer, S. Pattanayak, J. F. Joanny, L. Blanchoin, M. Théry, Force balance of opposing diffusive motors generates polarity-sorted microtubule patterns. *Proc. Natl. Acad. Sci. U. S. A.* **121** (2024).
9. S. Pattanayak, A. Sciortino, L. Blanchoin, M. Théry, J.-F. Joanny, Phase separation by polar active transport. arXiv arXiv:2604.01057 [Preprint] (2026). https://doi.org/10.48550/arXiv.2604.01057.
10. R. Grover, J. Fischer, F. W. Schwarz, W. J. Walter, P. Schwille, S. Diez, Transport efficiency of membrane-anchored kinesin-1 motors depends on motor density and diffusivity. *Proc. Natl. Acad. Sci. U. S. A.* **113**, E7185–E7193 (2016).

### Supplementary movies

**Movie S1 - Motor domains on an infinite bilayer.**
Time-lapse fluorescence imaging showing the self-organization of KIF5B (magenta) and NCD (cyan) into alternating motor domains on an unconfined supported lipid bilayer (SLB). Data are also shown in Fig. 1A.

**Movie S2 - Motor domains on micropatterns.**
Time-lapse fluorescence imaging showing the self-organization of KIF5B (magenta) and NCD (cyan) into spatially segregated motor domains within a circular micropatterned SLB. Data are also shown in Fig. 1B.

**Movie S3 - Micropatterned area and motor ratio.**
Time-lapse fluorescence imaging showing the formation of KIF5B (magenta) and NCD (cyan) domains within circular micropatterned SLBs of decreasing areas (8000, 2000, 500, and 250 $\mu m^2$). Reducing the confinement area decreases the number of motor domains. Data are also shown in Fig. 1C.

**Movie S4 - Motor ratios on large micropattern.**
Time-lapse fluorescence imaging showing the self-organization of KIF5B (magenta) and NCD (cyan) within large circular confinements at varying NCD/KIF5B concentration ratios. Changes in the relative motor concentrations alter the number, size, and spatial distribution of motor domains. Data are also shown in Fig. 1C.

**Movie S5- Motor ratios on small micropattern.**
Time-lapse fluorescence imaging showing the formation of KIF5B (magenta) and NCD (cyan) domains within small circular confinements at varying motor concentration ratios. Changes in motor composition favor either a polar organization with a single peripheral NCD domain or a radially symmetric organization with multiple peripheral KIF5B domains. Data are also shown in Fig. 1C and Fig. 2A.

**Movie S6 - Low concentration of NCD on small micropatterns.**
Time-lapse fluorescence imaging showing the formation of a single peripheral NCD domain (cyan) and a KIF5B-enriched domain (magenta), resulting in a polar organization within a circular confinement. Data are also shown in Fig. 2A, B.

**Movie S7 - Low concentration of KIF5B on small micropatterns.**
Time-lapse fluorescence imaging showing the formation of multiple peripheral KIF5B domains (magenta) surrounding a central NCD-enriched region (cyan), resulting in a radially symmetric organization. Data are also shown in Fig. 2A, C.

**Movie S8. Simulations of motor-domain organization at varying confinement sizes and motor ratios.**
Time-lapse simulations showing the self-organization of opposing motors into spatially segregated domains within circular confinements ($R=10\mu m$) and various NCD/KIF5B ratios. The simulations reproduce the emergence of distinct motor-domain organizations observed experimentally, as well as the non-mirror outcome of low amount of NCD or KIF5B when distinct strength are given to each motors

**Movie S9 - NCD addition on a polar array.**
Time-lapse fluorescence imaging showing the reorganization of KIF5B (magenta) and NCD (cyan) domains following the addition of NCD to a pre-existing polar array. Increasing NCD concentration reverses the relative positions of the motor domains while maintaining the polar organization. Data are also shown in Fig. 2D.

**Movie S10 - KIF5B addition on a radial array.**
Time-lapse fluorescence imaging showing the reorganization of KIF5B (magenta) and NCD (cyan) domains following the addition of KIF5B to a pre-existing radial array. Increasing KIF5B concentration reverses the radial distribution of motor domains while maintaining a radially symmetric organization. Data are also shown in Fig. 2F.

**Movie S11 - MT density on large micropatterns.**
Time-lapse fluorescence imaging showing the self-organization of KIF5B (magenta) and NCD (cyan) within circular confinements of 2000 $\mu m^2$ at increasing microtubule concentrations (50, 200, 400, and 800 nM). Increasing microtubule density promotes the formation of additional motor domains and longer interfaces. Data are also shown in Fig. 3A.

**Movie S12 - MT density on small micropatterns.**
Time-lapse fluorescence imaging showing the self-organization of KIF5B (magenta) and NCD (cyan) within circular confinements of 500 $\mu m^2$ at increasing microtubule concentrations (100, 200, 400, and 800 nM). Increasing microtubule density promotes the formation of motor-domain interfaces, while the number of domains remains constrained by the confinement size. Data are also shown in Fig. 3C.

**Movie S13 - MT elongation on unconfined lipid bilayer.**
Time-lapse fluorescence imaging showing microtubule elongation following the addition of free tubulin on an unconfined SLB. Elongating microtubules disrupt the pre-existing force balance, leading to the destabilization and dissolution of motor domains. Images show microtubules (green), NCD (cyan), and KIF5B (magenta). Data are also shown in Fig. S2A.

**Movie S14 - Control without MT elongation on large micropatterns.**
Time-lapse fluorescence imaging showing the organization of KIF5B (magenta) and NCD (cyan) domains within circular confinements of 2000 $\mu m^2$ without the addition of free tubulin. The pre-existing motor domains remain organized over time in the absence of microtubule elongation. Data are also shown in Fig. S2C.

**Movie S15 - MT elongation on large micropatterns.**
Time-lapse fluorescence imaging showing the reorganization of KIF5B (magenta) and NCD (cyan) domains following the addition of free tubulin within circular confinements of 2000 $\mu m^2$. Microtubule elongation reduces the number of motor domains and their interface length, leading to the formation of a bipolar organization. Data are also shown in Fig. 3E.

**Movie S16 - MT elongation on small micropatterns.**
Time-lapse fluorescence imaging showing the reorganization of KIF5B (magenta) and NCD (cyan) domains following the addition of free tubulin within circular confinements of 500 $\mu m^2$. Microtubule elongation reduces the number of domains and their interface length, resulting in a polar organization. Data are also shown in Fig. 3G.

**Movie S17 - Aspect ratio of large micropatterns.**
Time-lapse fluorescence imaging showing the self-organization of KIF5B (magenta) and NCD (cyan) within micropatterned SLBs of fixed area (500 µm²) and increasing aspect ratio. Increasing confinement elongation promotes the formation of two peripheral NCD domains separated by a central KIF5B domain, resulting in bipolar organization. Data are also shown in Fig. 4A.

**Movie S18 - Aspect ratio of small micropatterns.**
Time-lapse fluorescence imaging showing the self-organization of KIF5B (magenta) and NCD (cyan) within micropatterned SLBs of fixed area (2000 µm²) and increasing aspect ratio. Increasing confinement elongation reduces the number of peripheral NCD domains and favors the formation of a bipolar organization. Data are also shown in Fig. 4A.

**Movie S19. Motor ratio on elongated ellipses.**
Time-lapse fluorescence imaging showing the self-organization of KIF5B (magenta) and NCD (cyan) within elongated elliptical micropatterned supported lipid bilayers (SLBs) of 2000 µm² (Aspect ratio of 60) at varying NCD/KIF5B concentrations (10/10, 20/10, 30/10, 30/5, 50/5, and 50/2.5 nM). Changes in the relative motor concentrations regulate the number and size of motor domains, leading to the formation of alternating polarity-sorted domains along the elongated confinement. Data are also shown in Fig. 4C.

**Movie S20. Variation of bar length with 30 nM NCD and 10nM KIF5B**
Time-lapse fluorescence imaging showing the self-organization of KIF5B (magenta) and NCD (cyan) within elongated micropatterned SLBs of increasing lengths (6, 12, 22, 43, 65, and 109 µm) at fixed motor concentrations of 30 nM NCD and 10 nM KIF5B. Increasing confinement length promotes the formation of additional motor domains. Data are also shown in Fig. 4F.

**Movie S21 - Addition of Taxol on PtK2 cells expressing GFP-tubulin.**
Time-lapse fluorescence imaging showing the reorganization of MTs in PtK2 cells expressing GFP-tubulin upon treatment with 5µM Taxol. Images were taken every 30 minutes. Scale bar correspond to 20 µm.
Data are also shown in Fig. 5A.

**Movie S22 - Washout of Nocodazole and addition of Taxol on PtK2 cells expressing GFP-tubulin.**
Time-lapse fluorescence imaging showing the reorganization of MTs in PtK2 cells expressing GFP-tubulin upon washout of Nocodazole after overnight treatment and synchronous addition of 5µM Taxol. Images were taken every 1 minutes. Scale bar correspond to 20 µm.
Data are also shown in Fig. 5B.

**Movie S23 - Washout of Nocodazole and addition of Taxol on PtK2 expressing GFP-tubulin and micropatterned on circles**
Time-lapse fluorescence imaging showing the reorganization of MTs in PtK2 cells expressing GFP-tubulin upon washout of Nocodazole after overnight treatment and synchronous addition of 5µM Taxol. Cells were plated on micropatterned discs. Images were taken every 10 minutes. Scale bar correspond to 20 µm.
Data are also shown in Fig. 5E and Sup. Fig. S4A).

**Movie S24 - Washout of Nocodazole and addition of Taxol on PtK2 expressing GFP-tubulin and micropatterned on 12-micron-wide lines**

Time-lapse fluorescence imaging showing the reorganization of MTs in PtK2 cells expressing GFP-tubulin upon washout of Nocodazole after overnight treatment and synchronous addition of 5µM Taxol. Cells were plated on micropatterned 12-micron-wide lines. Images were taken every 10 minutes. Scale bar correspond to 20 µm.

Data are also shown in Fig. 5F and Sup. Fig. S4B).
